# Multiplex, quantitative, high-resolution imaging of protein:protein complexes via hybridization chain reaction

**DOI:** 10.1101/2023.07.22.550181

**Authors:** Samuel J. Schulte, Boyoung Shin, Ellen V. Rothenberg, Niles A. Pierce

**Author notes:** **Corresponding Author Niles A. Pierce** – Division of Biology & Biological Engineering, California Institute of Technology, Pasadena, California 91125, United States; Division of Engineering & Applied Science, California Institute of Technology, Pasadena, California 91125, United States.

## Abstract

Signal amplification based on the mechanism of hybridization chain reaction (HCR) facilitates spatial exploration of gene regulatory networks by enabling multiplex, quantitative, high-resolution imaging of RNA and protein targets. Here, we extend these capabilities to the imaging of protein:protein complexes, using proximity-dependent cooperative probes to conditionally generate a single amplified signal if and only if two target proteins are colocalized within the sample. HCR probes and amplifiers combine to provide automatic background suppression throughout the protocol, ensuring that even if reagents bind nonspecifically in the sample, they will not generate amplified background. We demonstrate protein:protein imaging with high signal-to-background in human cells, mouse proT cells, and highly autofluorescent formalin-fixed paraffin-embedded (FFPE) human breast tissue sections. Further, we demonstrate multiplex imaging of 3 different protein:protein complexes simultaneously and validate that HCR enables accurate and precise relative quantitation of protein:protein complexes with subcellular resolution in an anatomical context. Moreover, we establish a unified framework for simultaneous multiplex, quantitative, high-resolution imaging of RNA, protein, and protein:protein targets, with 1-step, isothermal, enzyme-free HCR signal amplification performed for all target classes simultaneously.

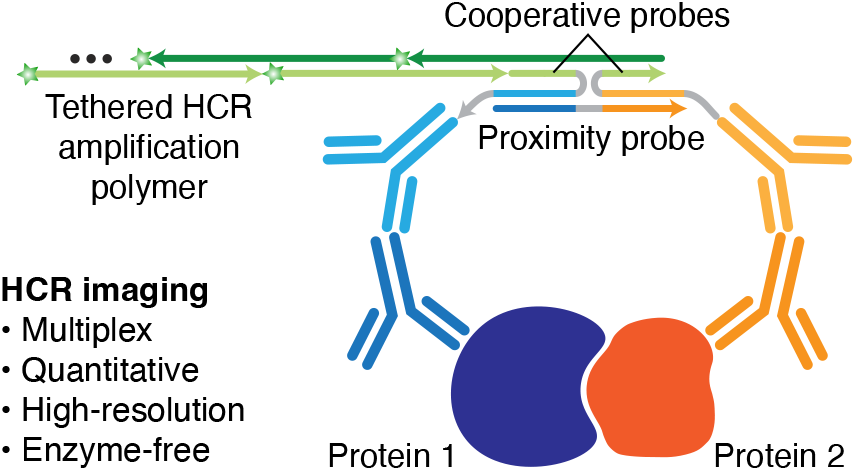

## INTRODUCTION

Methods for imaging molecular complexes^1, 2^ have been comparatively less explored than methods for imaging RNA and protein targets,^3–8^ yet represent an important frontier for spatial exploration of the interactome. Generating one signal conditional on the proximity of two molecules provides a sub-diffraction-limit readout, in contrast to independent imaging of the same two molecules with two signals. Protein:protein complexes play central roles in diverse cellular processes including transcription, translation, signaling, development, and disease.^9–12^ To date, imaging of protein:protein complexes has predominantly been performed using proximity ligation assays (PLA) that exploit enzyme-mediated ligation and rolling circle amplification,^13–17^ leading to challenges with both false-negatives (formation of non-circular ligation products^18^) and false-positives (background evident in technical controls that omit one reaction component^17^), as well as issues with cost and variable enzyme activity.^13, 17^ Alternatively, to avoid the use of enzymes, a proximity-based HCR approach has been developed that uses a kinetic trigger mechanism to desequester an HCR initiator if two probes are bound to proximal target proteins;^18, 19^ this approach has so far been limited to 1-plex applications.

Over the course of nearly two decades, we have developed simple and robust HCR RNA in situ hybridization (RNA-ISH) and immunohistochemistry (IHC) methods that enable biologists, drug developers, and pathologists to perform multiplex, quantitative, high-resolution imaging of RNA and protein targets in highly autofluorescent samples.^20–26^ Here, we sought to use HCR principles to extend these benefits to the imaging of protein:protein complexes. An HCR amplifier consists of two species of kinetically trapped DNA hairpins (h1 and h2) that co-exist metastably in solution, storing the en-ergy to drive conditional self-assembly of an HCR amplification polymer upon exposure to a cognate initiator sequence (i1; Figure 1A).^27^ Using HCR RNA-ISH, an RNA target is detected using one or more pairs of split-initiator DNA probes, each carrying a fraction of HCR initiator i1 (Figure 1B).^24^ Probe pairs that hybridize specifically to proximal binding sites on the target RNA colocalize a full HCR initiator i1 capable of triggering HCR signal amplification. Meanwhile, any individual probes that bind nonspecifically in the sample do not colocalize full HCR initiator i1 and do not trigger HCR. Using HCR IHC, a protein target is detected using an unlabeled primary antibody probe, which in turn is detected by an initiator-labeled secondary antibody probe that carries an HCR initiator i1 capable of triggering HCR signal amplification (Figure 1C).^26^

**Figure 1.**
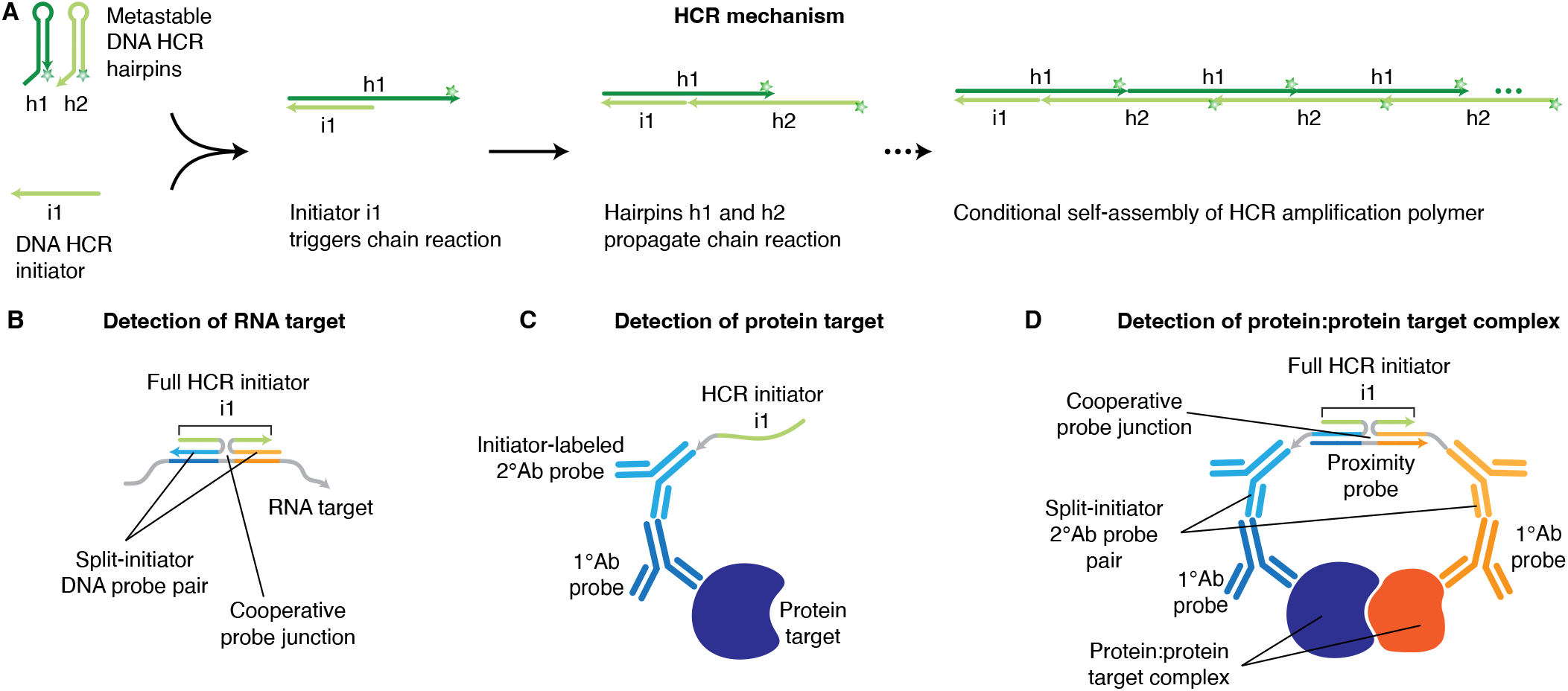
Applying HCR principles to enable simple and robust imaging of protein:protein complexes. (A) HCR mechanism.^27^ Stars denote fluorophores. Arrowhead indicates 3 end of each strand. (B) HCR RNA-ISH: an RNA target is detected using a pair of split-initiator DNA probes, each carrying a fraction of HCR initiator i1. (C) HCR IHC: a protein target is detected using an unlabeled primary antibody probe and an initiator-labeled secondary antibody probe carrying HCR initiator i1. (D) HCR protein:protein imaging: a protein:protein target complex is detected with a pair of unlabeled primary antibodies, a pair of split-initiator secondary antibodies each carrying a fraction of HCR initiator i1, and a proximity probe.

We hypothesized that the split-initiator concept from HCR RNA-ISH (Figure 1B) could be generalized using the antibody probes of HCR IHC (Figure 1C) to enable simple and robust HCR imaging of protein:protein complexes using a split-initiator antibody probe pair in conjunction with a new proximity probe (Figure 1D). Here, we demonstrate that this combination of proximity-dependent cooperative probes and metastable HCR amplifiers enables multiplex, quantitative, high-resolution imaging of protein:protein complexes, including full compatibility with HCR RNA-ISH and HCR IHC.

## RESULTS AND DISCUSSION

### HCR imaging of protein:protein complexes using a 3-stage protocol

HCR imaging of protein:protein complexes is performed using the 3-stage protocol summarized in Figure 2A. In the detection stage, two protein targets are detected with unlabeled primary antibody probes that are in turn detected by a pair of split-initiator secondary antibody probes (p1 and p2) each carrying a fraction of HCR initiator i1 and a proximity domain. In the proximity stage, if the two protein targets are colocalized in the sample, the proximity probe is able to hybridize to p1 and p2 to colocalize a full HCR initiator i1 capable of triggering HCR signal amplification. Note that the proximity probe creates a cooperative probe junction (Figure 1D) inspired by the cooperative probe junction cre-ated in HCR RNA-ISH (Figure 1B), with the DNA proximity probe taking the place of the RNA target. Any split-initiator probes that bind nonspecifically or to isolated protein targets in the sample can hybridize to the proximity probe, but will not colocalize a full HCR initiator i1 and will not trigger HCR. In the amplification stage, each colocalized full HCR initiator i1 triggers self-assembly of metastable fluorophore-labeled HCR hairpins (h1 and h2) into a tethered fluorescent amplification polymer to generate an amplified signal at the site of the protein:protein target complex.

**Figure 2.**
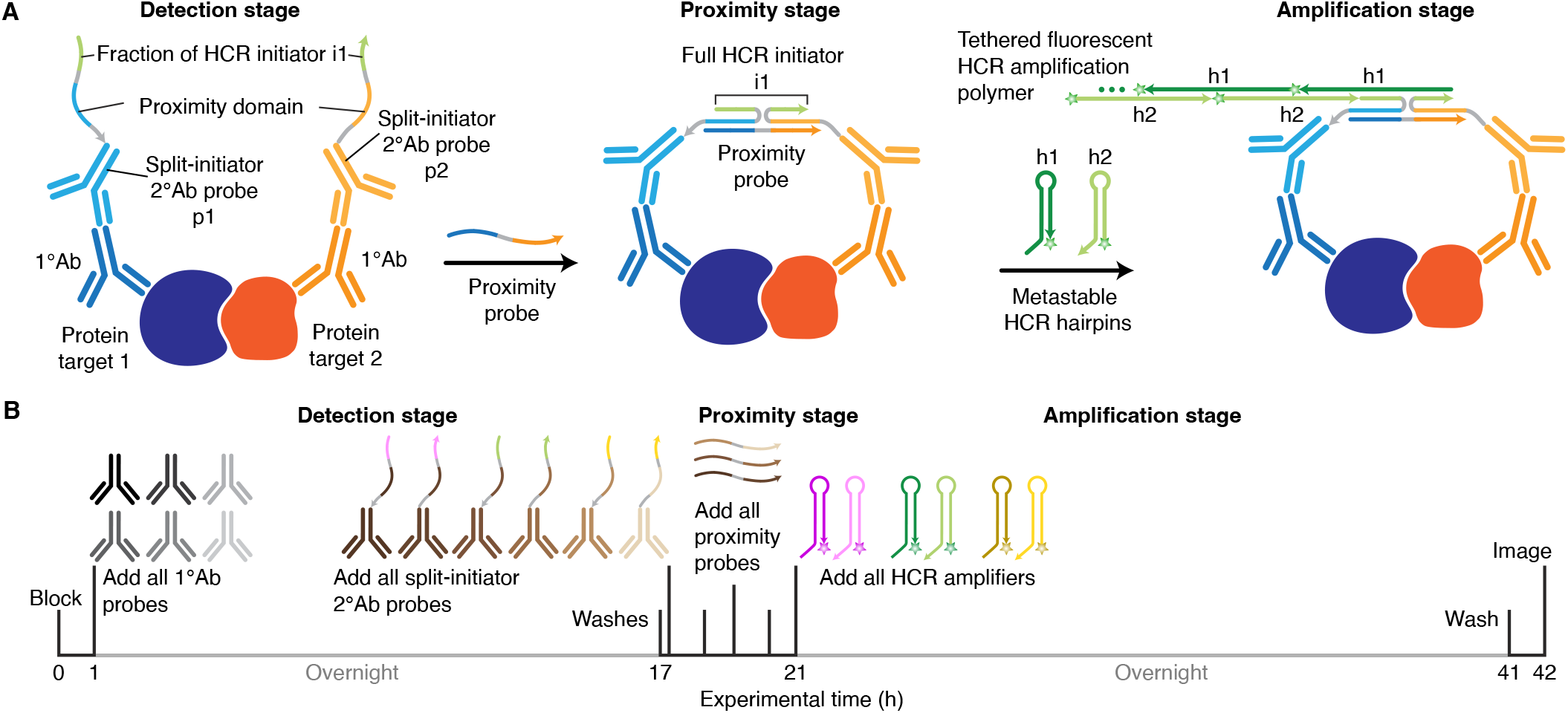
Imaging protein:protein complexes using HCR. (A) Three-stage protocol. Detection stage: unlabeled primary antibody probes bind to protein targets 1 and 2; wash; split-initiator secondary antibody probes p1 and p2 bind to primary antibody probes; wash. Proximity stage: if p1 and p2 are proximal, a proximity probe hybridizes to the proximity domains of p1 and p2 to colocalize full HCR initiator i1. Amplification stage: colocalized full HCR initiator i1 triggers self-assembly of fluorophore-labeled HCR hairpins into a tethered fluorescent amplification polymer; wash. (B) Multiplexing timeline. The same three-stage protocol is used independent of the number of protein:protein target complexes.

### Imaging protein:protein complexes in human cells, mouse proT cells, and FFPE human breast tissue sections

To evaluate the performance of our split-initiator approach for imaging protein:protein complexes, we compared the fluorescence intensity between three pairs of biological sample types using the same imaging settings for both sample types. Positive samples are expected to form the protein:protein complex of interest; negative samples are expected to have minimal or no formation of the protein:protein complex of interest. For each pair of sample types, we calculate an estimated signal-to-background ratio, using the positive sample type to estimate signal plus background and the negative sample type to estimate background. This approach yields a conservative estimate of performance, as characterizing background in a sample containing little or no protein:protein target complex places an upper bound on background and hence a lower bound on signal-to-background.

First, we compared the fluorescence intensity for the *β*-catenin:E-cadherin complex in A-431 and HeLa adherent human cell lines. While A-431 cells form the *β*-catenin:E-cadherin complex at the cell membrane of intercellular junctions,^28^ HeLa cells express N-cadherin rather than E-cadherin^29, 30^ and therefore lack the *β*-catenin:E-cadherin complex. As expected, A-431 cells (Figure 3A) display strong signal at intercellular junctions and HeLa cells display no visible staining (Figure 3B), with a signal-to-background ratio of 26 ± 4 between the two cell lines (mean ± SEM for representative regions of *N* = 3 replicate wells on a slide).

**Figure 3.**
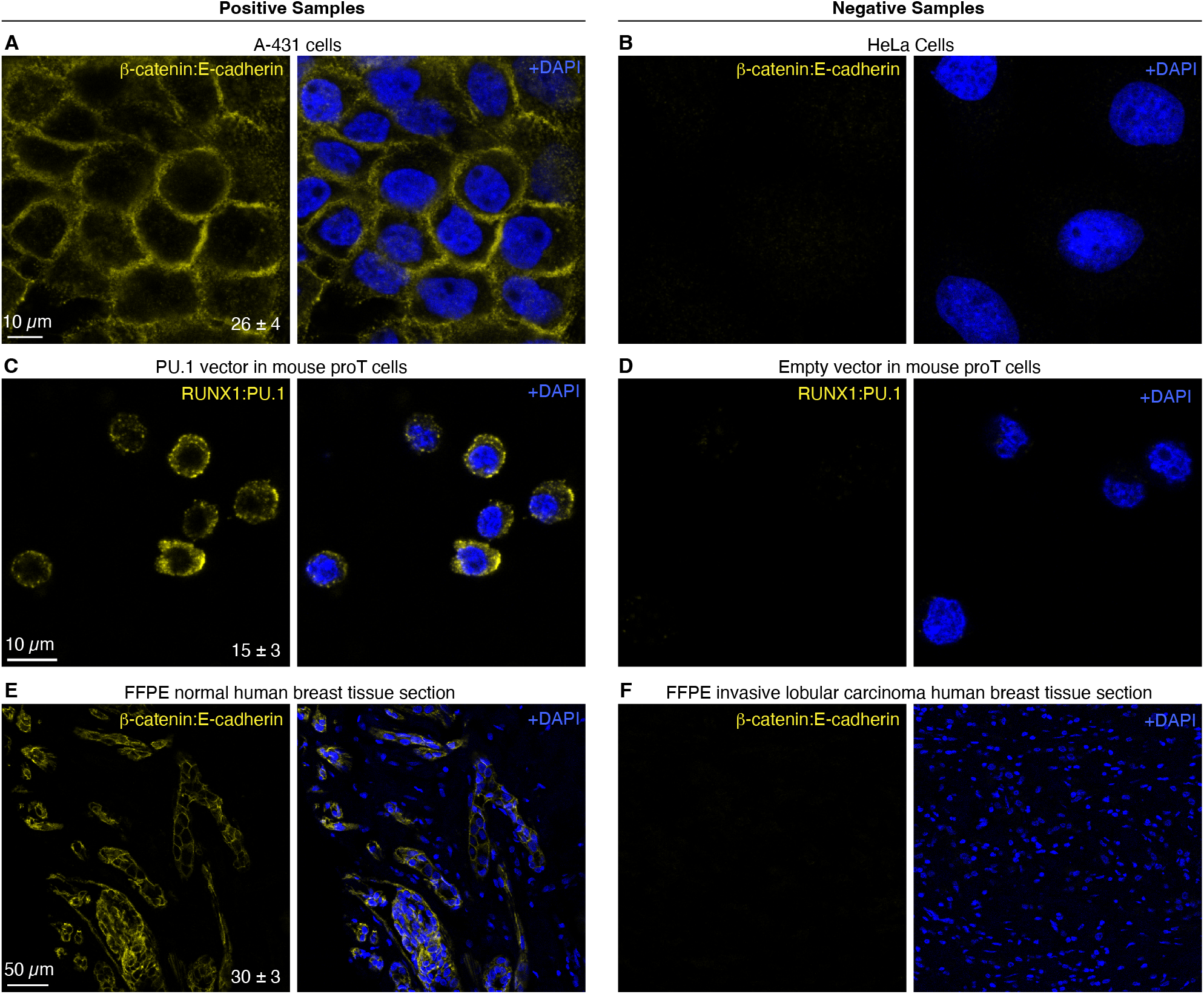
Imaging protein:protein complexes in human cells, mouse proT cells, and FFPE human breast tissue sections. (A,B) Imaging *β*-catenin:E-cadherin target complex in A-431 cells expressing *β*-catenin and E-cadherin (panel A) or HeLa cells expressing N-cadherin instead of E-cadherin (panel B). (C,D) Imaging RUNX1:PU.1 target complex in Scid.adh.2C2 mouse proT cells retrovirally transduced with a PU.1-expressing vector (panel C) or an empty vector (panel D). (E,F) Imaging *β*-catenin:E-cadherin target complex in 5 *μ*m FFPE human breast tissue sections from the same patient: normal (panel E) or invasive lobular carcinoma (panel F). All panels: confocal image; single optical section; 0.18*×*0.18*×*0.8 *μ*m pixels (panels A-D) or 0.57*×*0.57*×*3.3 *μ*m pixels (panels E,F). Signal-to-backround ratio for each row (mean ± SEM for representative regions of *N* = 3 replicate samples). See Sections S2.2–S2.4 for additional data.

Next, we imaged Scid.adh.2C2^31^ mouse proT cells in search of the RUNX1:PU.1 target complex. The Scid.adh.2C2 cell line has emerged as a useful proT cell line for studying T cell development, with exogenous introduction of PU.1 protein capable of reverting the cell line to an earlier developmental time point, in part via direct or indirect interactions between PU.1 and other proteins such as RUNX1.^32–34^ Because the Scid.adh.2C2 cell line does not endogenously express the PU.1 protein,^31^ Scid.adh.2C2 cells cannot natively form the RUNX1:PU.1 complex. When the Scid.adh.2C2 cell line is retrovirally transduced with PU.1, it is unknown whether PU.1 forms a complex with RUNX1 or interacts less directly.^33^ Here, imaging the RUNX1:PU.1 target complex, we observe signal in cells retrovirally transduced with a PU.1-containing vector (Figure 3C) and no visible staining for cells retrovirally transduced with an empty vector (Figure 3D), with a signal-to-background ratio of 15 ± 3 between the two experiment types (mean ± SEM for representative regions of *N* = 3 replicate wells on a slide). These results provide evidence that RUNX1 and PU.1 are spatially colocalized in Scid.adh.2C2 cells and not merely logically linked.

To test performance in highly autofluorescent samples, we detected the *β*-catenin:E-cadherin complex in normal and pathological FFPE human breast tissue sections. The *β*-catenin:E-cadherin complex is robustly formed in normal breast epithelial cells, but the expression of and interaction between the *β*-catenin and E-cadherin proteins is interrupted when breast epithelial cells become cancerous in the invasive lobular carcinoma disease process.^35, 36^ We obtained paired normal and invasive lobular carcinoma FFPE breast tissue sections from the same patient and evaluated them for the *β*-catenin:E-cadherin complex, observing strong signal in normal breast tissue (Figure 3E) and no visible staining in cancerous tissue (Figure 3F), with a signal-to-background ratio of 30 ± 3 between the two tissue types (mean ± SEM for representative regions of *N* = 3 replicate sections).

In summary, protein:protein complexes are imaged with high signal-to-background across three different paired sample types, including highly autofluorescent FFPE tissues.

### Multiplex protein:protein imaging

HCR RNA-ISH and HCR IHC enable straightforward multiplexing for RNA and protein targets to allow multidimensional analyses of gene expression in an anatomical context.^20–22, 24–26^ To likewise enable multiplex imaging of protein:protein complexes, we used NUPACK^37, 38^ to design proximity probes for three orthogonal HCR amplifiers. Figure 4 demonstrates multiplex protein:protein imaging for three target complexes that localize to different compartments of A-431 adherent human cells: cytoskeletal *α*-tubulin:*β*-tubulin complex, membranous *β*-catenin:E-cadherin complex, and nuclear speckle SC35:SON complex. High signal-to-background is observed for all three protein:protein target complexes, with background estimated based on technical control experiments that omit the primary and secondary antibody probes for one protein or the other within a given complex (see Table S13 for details). Multiplexing is straightforward using a three-stage protocol inde-pendent of the number of protein:protein target complexes (Figure 2B): all protein targets are detected in parallel, proximity is verified for all protein target pairs in parallel, and amplification is performed for all colocalized full HCR initiators in parallel.

**Figure 4.**
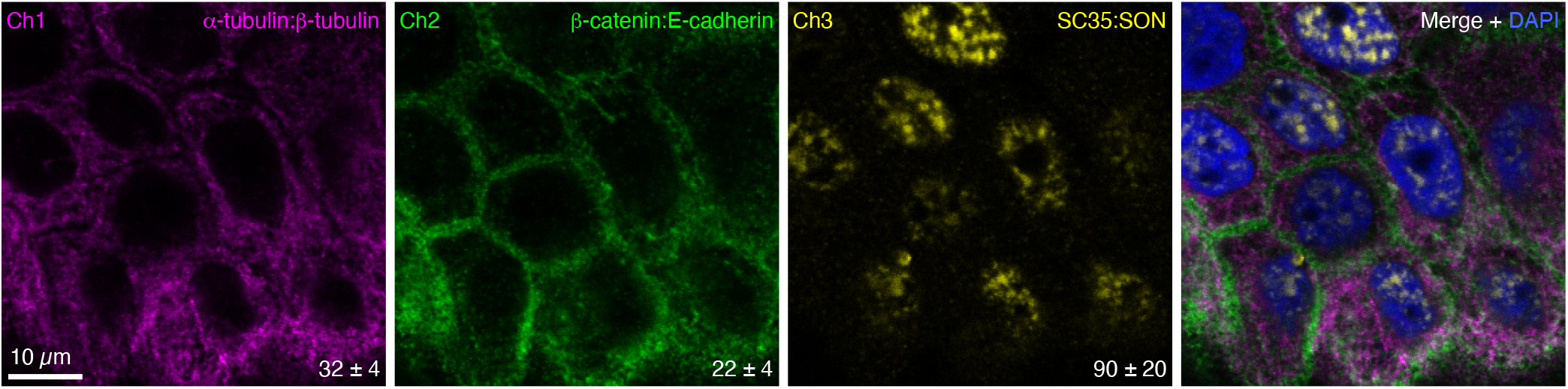
Multiplex imaging of protein:protein complexes. Three-channel confocal image in A-431 cells; single optical section; 0.18*×*0.18*×*0.8 *μ*m pixels. Ch1: cytoskeletal *α*-tubulin:*β*-tubulin complex (Alexa488). Ch2: membranous *β*-catenin:E-cadherin complex (Alexa546). Ch3: nuclear speckle SC35:SON complex (Alexa647). Signal-to-background ratio for each channel (mean ± SEM for representative regions of *N* = 3 replicate wells on a slide). See Section S2.5 for additional data.

### qHCR imaging: relative quantitation of protein:protein complexes with subcellular resolution

We have previously demonstrated that HCR imaging enables accurate and precise relative quantitation of both RNA and protein targets with subcellular resolution in an anatomical context, generating an amplified signal that scales approximately linearly with the number of target molecules per imaging voxel.^24–26^ Here, we validate that the proximity probe and split-initiator antibody probe pair preserve the quantitative nature of HCR imaging for protein:protein target complexes. To test relative quantitation, we detect each protein in the complex with an unlabeled primary antibody probe as usual, and then redundantly detect each primary antibody probe with two batches of split-initiator secondary antibody probes, where each batch interacts with a different proximity probe and triggers a different spectrally distinct HCR amplifier (Figure 5A), yielding a two-channel image (Figure 5B). If HCR signal scales approximately linearly with the number of target protein:protein complexes per voxel, a two-channel scatter plot of normalized voxel intensities will yield a tight linear distribution with zero intercept.^25^ Consistent with expectation, we observe high accuracy (linearity with zero intercept) and precision (scatter around the line) for subcellular voxels in both cultured human cells (Figure 5C; top) and highly autofluorescent FFPE human breast tissue (Figure 5C; bottom).

**Figure 5.**
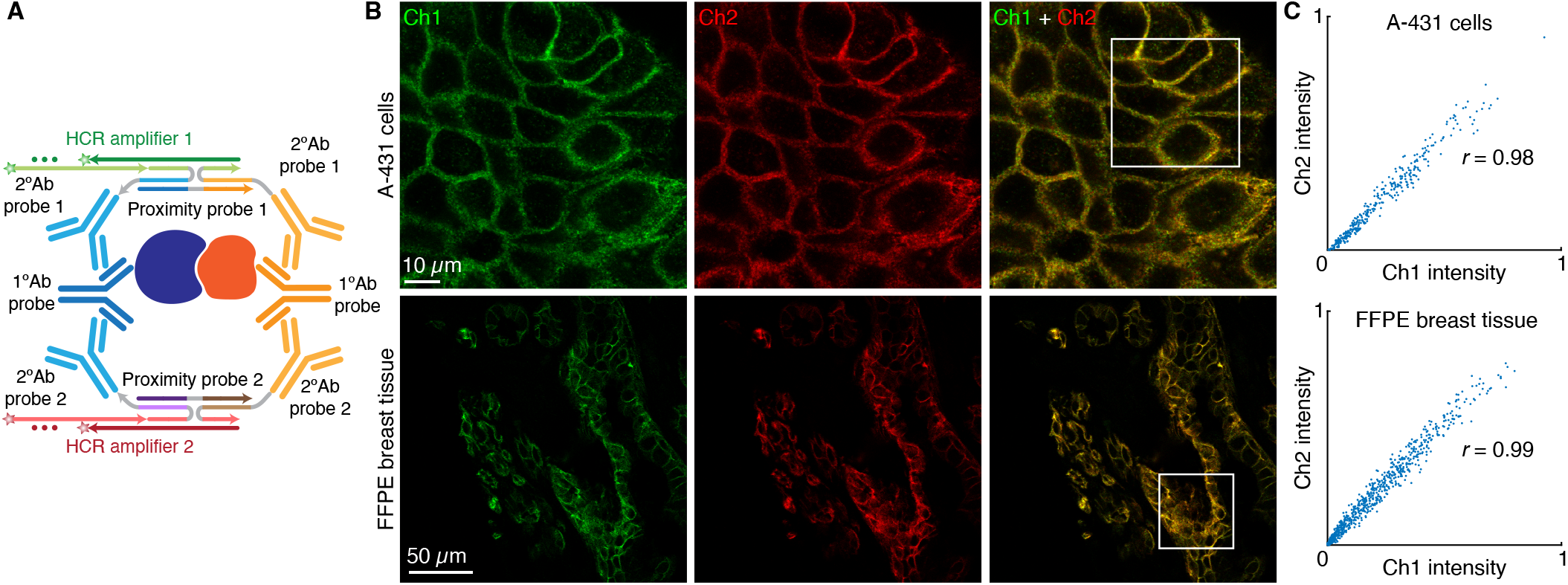
qHCR imaging: relative quantitation of protein:protein complexes with subcellular resolution in an anatomical context. (A) Two-channel redundant detection of a protein:protein complex: each target protein is detected by an unlabeled primary antibody probe and two batches of secondary antibody probes that interact with orthogonal proximity probes to colocalize full HCR initiators that trigger orthogonal spectrally distinct HCR amplifiers (Ch1: Alexa546; Ch2: Alexa647). (B) Two-channel confocal images. Top: *β*-catenin:E-cadherin complex in A-431 cells (0.18*×*0.18*×*0.8 *μ*m pixels). Bottom: *β*-catenin:E-cadherin complex in a 5 *μ*m FFPE normal human breast tissue section (0.57*×*0.57*×*3.3 *μ*m pixels). (C) High accuracy and precision for protein:protein relative quantitation in an anatomical context. Highly correlated normalized signal (Pearson correlation coefficient, *r*) for subcellular voxels in the indicated regions in panel B. Top: 2.0*×*2.0*×*0.8 *μ*m voxels. Bottom: 2.0*×*2.0*×*3.3 *μ*m voxels. Accuracy: linearity with zero intercept. Precision: scatter around the line. See Section S2.6 for additional data.

### Simultaneous multiplex imaging of protein, protein:protein, and RNA targets

We have previously shown that HCR RNA-ISH and HCR IHC enable multiplex, quantitative, high-resolution RNA and protein imaging in highly autofluorescent samples.^26^ Here, we demonstrate compatible multiplex imaging of protein, protein:protein, and RNA targets using initiator-labeled antibody probes for protein targets, proximity probes and split-initiator antibody probe pairs for protein:protein targets, and split-initiator DNA probe pairs for RNA targets, with simultaneous HCR signal amplification for all target classes (Figure 6A). In A-431 adherent human cells, mitochondrial HSP60 protein targets, cytoskeletal *α*-tubulin:*β*-tubulin protein:protein target complexes, and nuclear *U6* RNA targets are all imaged simultaneously (Figure 6B) with high signal-to-background (see Table S18 for additional details).

**Figure 6.**
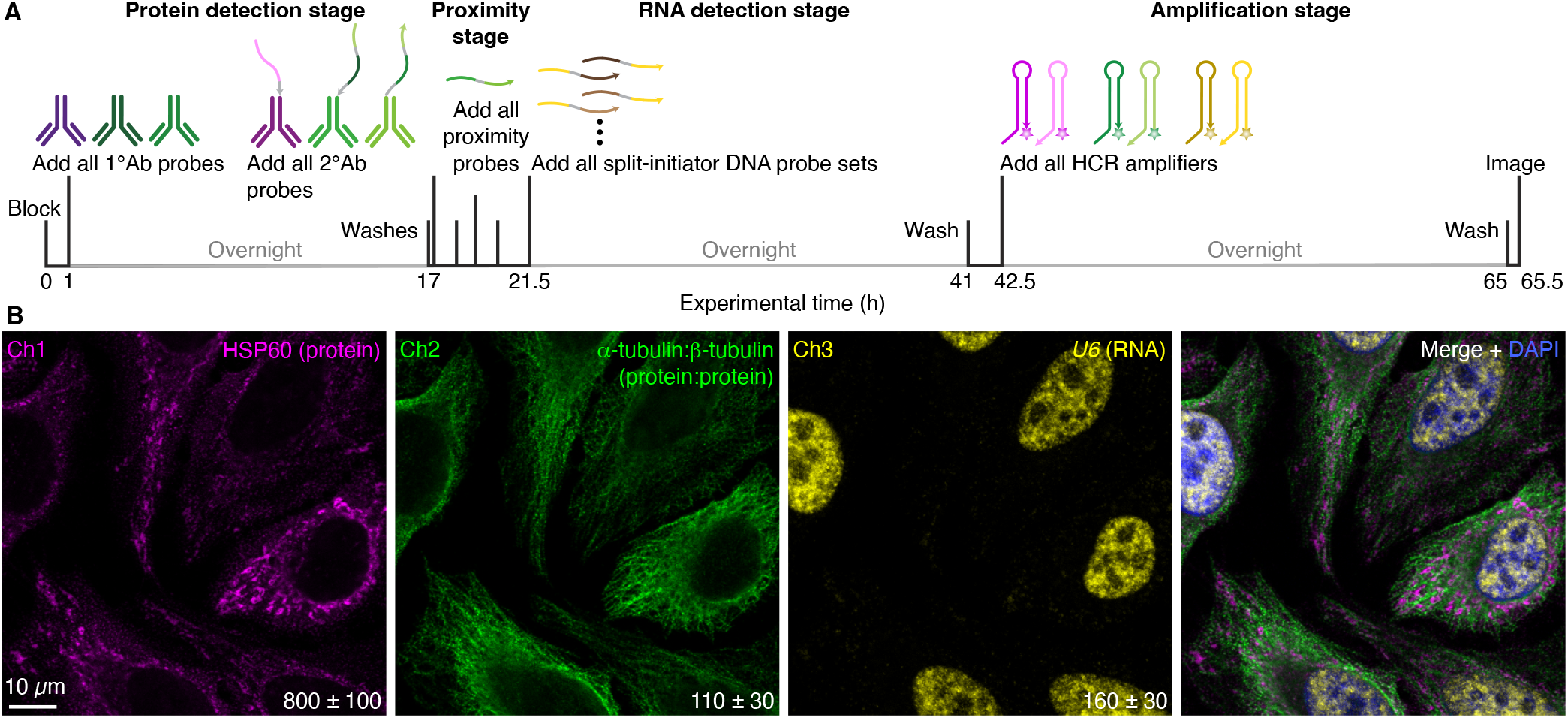
Simultaneous multiplex protein, protein:protein, and RNA imaging using HCR. (A) Four-stage protocol. Protein detection stage: unlabeled primary antibody probes bind to protein targets; wash; secondary antibody probes bind to primary antibody probes (initiator-labeled 2°Ab probes associated with individual protein targets carry full initiators; split-initiator 2°Ab probes associated with protein:protein target complexes carry a fraction of HCR initiator i1 and a proximity domain); wash. Proximity stage: proximity probes colocalize a full HCR initiator i1 for protein:protein target complexes; wash. RNA detection stage: split-initiator DNA probes bind to RNA targets; wash. Amplification stage: initiators trigger self-assembly of fluorophore-labeled HCR hairpins into tethered fluorescent HCR amplification polymers; wash. (B) Three-channel confocal image in HeLa cells; single optical section; 0.18*×*0.18*×*0.8 *μ*m pixels. Ch1: mitochondrial HSP60 protein (Alexa488). Ch2: cytoskeletal *α*-tubulin:*β*-tubulin complex (Alexa546). Ch3: nuclear *U6* RNA (Alexa647). Signal-to-background ratio for each channel (mean ± SEM for representative regions of *N* = 3 replicate wells on a slide). See Section S2.7 for additional data.

### Unified framework for multiplex, quantitative, high-resolution imaging

We have shown that HCR imaging provides a unified framework for multiplex, quantitative, high-resolution imaging of RNA targets, protein targets, and pro-tein:protein target complexes simultaneously. High signal-to-background is achieved even in highly autofluorescent samples. As a natural property of this method, the amplified signal scales approximately linearly with target abundance, enabling accurate and precise relative quantitation of each target with subcellular resolution in an anatomical context. Using the validated proximity probes presented here, up to three protein:protein target complexes can be imaged simultaneously, in combination with RNA and/or protein targets of choice.

### Automatic background suppression throughout the protocol

As is the case for HCR RNA-ISH, the use of split-initiator probes during the detection stage and metastable HCR hairpins during the amplification stage provides automatic background suppression throughout the protocol, ensuring that even if reagents bind nonspecifically in the sample, they do not generate amplified background.^24^ During the detection stage, any individual probes that bind nonspecifically in the sample do not colocalize a full HCR initiator and do not trigger HCR. Likewise, during the proximity stage for protein:protein imaging, any proximity probes that bind nonspecifically in the sample lack the ability to initiate HCR. During the amplification stage, any HCR hairpins that bind non-specifically in the sample are kinetically trapped and do not trigger formation of an HCR amplification polymer. Automatic background suppression enhances signal-to-background and quantitative accuracy and precision.^24^

### Split-initiator primary antibody probes vs split-initiator secondary antibody probes

The work presented here employs unlabeled primary antibody probes and split-initiator secondary antibody probes to detect protein:protein complexes, thereby requiring that each primary antibody be of a different isotype or raised in a different host species. Given the large libraries of commercially available antibodies available to users, this requirement is often not an impediment. For example, to image three protein:protein complexes simultaneously (Figure 4), we employ chicken IgY, mouse IgG1, mouse IgG2a, mouse IgG2b, guinea pig IgG, and rabbit IgG primary antibody probes to detect the six target proteins. How-ever, when it is desirable to use multiple primary antibodies raised in the same host species or of the same isotype, just as HCR IHC can be performed using initiator-labeled primary antibody probes,^26^ there is the option to perform HCR protein:protein imaging using split-initiator primary antibody probes (see the diagrams of Figure S1). Because antibody-oligo conjugation can sometimes interfere with target recognition, there is a practical advantage to using split-initiator secondary antibody probes as we do here: using a small library of validated split-initiator secondary antibody probes, users can plug-and-play with large libraries of unmodified primary antibody probes with no need to validate antibody-oligo conjugation for each new target protein. Additionally, because multiple split-initiator secondary antibody probes can bind to each primary antibody probe, there is the potential for proximity probes to colocalize multiple full HCR initiators per target complex, triggering growth of multiple tethered fluorescent amplification polymers and increasing amplification gain.

### Simple, robust, enzyme-free imaging of protein:protein complexes

In conclusion, HCR principles^27^ drawn from the emerging discipline of dynamic nucleic acid nanotechnology lead to an enzyme-free approach for imaging protein:protein complexes that retains the desirable simplicity and robustness of RNA and protein imaging using HCR RNA-ISH^24^ and HCR IHC.^26^

## METHODS

### Probes, amplifiers, and buffers

Probes, amplifiers, and buffers were obtained from Molecular Technologies, a non-profit academic resource within the Beckman Institute at Caltech. Details on the probes, amplifiers, and buffers for each experiment are displayed in Table S1 for HCR imaging of protein:protein complexes, in Table S2 for HCR RNA-ISH, and in Table S3 for HCR IHC.

### HCR imaging of protein:protein complexes

HCR imaging of protein:protein complexes, with optional co-detection of protein and RNA targets, was performed in adherent human cell lines (A-431 or HeLa) using the protocol detailed in Section S1.8. A-431 cells (ATCC, CRL-1555) were cultured in Dulbecco’s Modified Eagle Medium (DMEM) with high glucose and pyruvate (Gibco, 11995-073) supplemented with 10% fetal bovine serum (FBS) (Sigma-Aldrich, F4135). HeLa cells (ATCC, CRM-CCL-2) were cultured in Eagle’s Minimum Essential Medium (EMEM) (ATCC, 30-2003) supplemented with 10% FBS (Sigma-Aldrich, F4135). HCR imaging of protein:protein complexes was performed in Scid.adh.2C2 mouse proT cells^31^ cultured in RPMI1640 media (Gibco, 31800022) supplemented with 10% FBS (Sigma-Aldrich, F2442), 1*×* Penicillin-Streptomycin-Glutamine (Gibco, 10378-016), 0.1 mM sodium pyruvate (Gibco, 11360-070), 1*×*MEM non-essential amino acids (Gibco, 11140-050), and 50 *μ*M *β*-mercaptoethanol (Gibco, 21985-023) using the protocol detailed in Section S1.9. HCR imaging of protein:protein complexes was performed in 5 *μ*m FFPE normal human breast tissue sections (Acepix Biosciences, HuN-06-0027) and 5 *μ*m FFPE invasive lobular carcinoma human breast tissue sections (Acepix Biosciences, HuC-06-0101) from the same patient using the protocol detailed in Section S1.10.

### Microscopy

Confocal microscopy was performed using a Leica Stellaris 8 inverted confocal microscope. All images are displayed without background subtraction. Each channel (except for DAPI) is displayed with 0.01% of pixels saturated across three replicates. Details on the objectives, excitation wavelengths, detectors, and detection wavelengths used for each experiment are displayed in Table S5.

### Image analysis

Image analysis was performed as detailed in Section S1.7, including: definition of raw pixel intensities; measurement of signal, background, and signal-to-background; and calculation of normalized subcellular voxel intensities for qHCR imaging.

## ASSOCIATED CONTENT

### Supporting Information

Materials, additional methods, and replicate data.

## AUTHOR INFORMATION

### Authors

**Samuel J. Schulte** – *Division of Biology & Biological Engineering, California Institute of Technology, Pasadena, California 91125, United States;*

**Boyoung Shin** – *Division of Biology & Biological Engineering, California Institute of Technology, Pasadena, California 91125, United States;*

**Ellen V. Rothenberg** – *Division of Biology & Biological Engineering, California Institute of Technology, Pasadena, California 91125, United States;*

### Notes

The authors declare competing financial interests in the form of patents, pending patent applications, and the startup company Molecular Instruments.

## ACKNOWLEDGMENTS

We thank M. E. Bronner for reading a draft of the manuscript and G. Shin of the Molecular Technologies resource within the Beckman Institute at Caltech for providing HCR reagents. This work was funded by the National Institutes of Health (NIBIB R01EB006192 and NIGMS training grant GM008042 to S.J.S.) and by the Beckman Institute at Caltech (Programmable Molecular Technology Center, PMTC). The Leica Stellaris 8 confocal microscope in the Biological Imaging Facility within the Beckman Institute at Caltech was purchased with support from Caltech and the following Caltech entities: the Beckman Institute, the Resnick Sustainability Institute, the Division of Biology & Biological Engineering, and the Merkin Institute for Translational Research.

## ABBREVIATIONS USED

FFPE: formalin-fixed paraffin-embedded
HCR: hybridization chain reaction
IHC: immunohistochemistry
ISH: in situ hybridization
PLA: proximity ligation assay
qHCR: quantitative hybridization chain reaction

