## Supplementary Information for "Multiplex, quantitative, high-resolution imaging of protein:protein complexes via hybridization chain reaction"

**References** **57**

### S1 Additional materials and methods

#### S1.1 Probe and amplifier details for HCR imaging of protein:protein complexes

| Species | Sample | Protein:protein complex | Unlabeled 1° Ab probe<br>Split-initiator 2° Ab probe | Dilution factor<br>Working concentration ($\mu$g/mL) | Supplier (catalog #) | HCR amplifier | Figures |
|---|---|---|---|---|---|---|---|
| *H. sapiens sapiens* | A-431 or HeLa cells | $\beta$-catenin:E-cadherin | 1° mAb mouse IgG1 anti-$\beta$-catenin | 1:100 | Abc (ab237983) | | |
| | | | 2° pAb goat anti-mouse IgG1-B1-p1 | 1 | MT (A9111-AB1) | | |
| | | | 1° mAb rabbit IgG anti-E-cadherin | 1:200 | CST (3195S) | | |
| | | | 2° pAb donkey anti-rabbit IgG-B1-p2 | 1 | MT (A9230-ZB1) | B1-Alexa647 | 3AB, S2, S3 |
| *M. musculus* | Scid.adh.2C2 ProT cells | RUNX1:PU.1 | 1° pAb sheep IgG anti-RUNX1 | 1:300 | AO (ABIN350811) | | |
| | | | 2° pAb donkey anti-sheep IgG-B1-p1 | 1 | MT (A9233-AB1) | | |
| | | | 1° mAb rabbit IgG anti-PU.1 | 1:500 | Abc (ab227835) | | |
| | | | 2° pAb donkey anti-rabbit IgG-B1-p2 | 1 | MT (A9230-ZB1) | B1-Alexa647 | 3CD, S4, S5 |
| *H. sapiens sapiens* | Normal or pathological FFPE breast tissue | $\beta$-catenin:E-cadherin | 1° mAb mouse IgG1 anti-$\beta$-catenin | 1:100 | Abc (ab237983) | | |
| | | | 2° pAb goat anti-mouse IgG1-B1-p1 | 1 | MT (A9111-AB1) | | |
| | | | 1° mAb rabbit IgG anti-E-cadherin | 1:200 | CST (3195S) | | |
| | | | 2° pAb donkey anti-rabbit IgG-B1-p2 | 1 | MT (A9230-ZB1) | B1-Alexa647 | 3EF, S6, S7 |
| *H. sapiens sapiens* | A-431 cells | $\alpha$-tubulin:$\beta$-tubulin | 1° mAb mouse IgG2a anti-$\beta$-tubulin | 1:50 | Inv (MA5-16308) | | |
| | | | 2° pAb goat anti-mouse IgG2a-B6-p1 | 1 | MT (A9112-AB6) | | |
| | | | 1° pAb guinea pig IgG anti-$\alpha$-tubulin | 1:500 | Sy (302 204) | | |
| | | | 2° pAb donkey anti-guinea pig IgG-B6-p2 | 1 | MT (A9231-ZB6) | B6-Alexa488 | 4, S8, S9 |
| | | $\beta$-catenin:E-cadherin | 1° pAb chicken IgY anti-$\beta$-catenin | 1:100 | Aves (BCAT-0020) | | |
| | | | 2° pAb donkey anti-chicken IgY-B1-p1 | 1 | MT (A9232-AB1) | | |
| | | | 1° mAb mouse IgG2b anti-E-cadherin | 1:100 | Pr (60335-1-IG) | | |
| | | | 2° pAb goat anti-mouse IgG2b-B1-p2 | 1 | MT (A9113-ZB1) | B1-Alexa546 | 4, S8, S10 |
| | | SC35:SON | 1° mAb mouse IgG1 anti-SC35 | 1:200 | Abc (ab11826) | | |
| | | | 2° pAb goat anti-mouse IgG1-B9-p1 | 1 | MT (A9111-AB9) | | |
| | | | 1° pAb rabbit IgG anti-SON | 1:100 | Atlas (HPA023535) | | |
| | | | 2° pAb donkey anti-rabbit IgG-B9-p2 | 1 | MT (A9230-ZB9) | B9-Alexa647 | 4, S8, S11 |
| *H. sapiens sapiens* | A-431 cells | $\beta$-catenin:E-cadherin | 1° mAb mouse IgG1 anti-$\beta$-catenin | 1:100 | Abc (ab237983) | | |
| | | | 1° mAb rabbit IgG anti-E-cadherin | 1:200 | CST (3195S) | | |
| | | | 2° pAb goat anti-mouse IgG1-B1-p1 | 1 | MT (A9111-AB1) | | |
| | | | 2° pAb donkey anti-rabbit IgG-B1-p2 | 1 | MT (A9230-ZB1) | B1-Alexa546 | 5, S12, S15 |
| | | | 2° pAb goat anti-mouse IgG1-B6-p1 | 1 | MT (A9111-AB6) | | |
| | | | 2° pAb donkey anti-rabbit IgG-B6-p2 | 1 | MT (A9230-ZB6) | B6-Alexa647 | 5, S12, S16 |
| *H. sapiens sapiens* | Normal FFPE breast tissue | $\beta$-catenin:E-cadherin | 1° mAb mouse IgG1 anti-$\beta$-catenin | 1:100 | Abc (ab237983) | | |
| | | | 1° mAb rabbit IgG anti-E-cadherin | 1:200 | CST (3195S) | | |
| | | | 2° pAb goat anti-mouse IgG1-B1-p1 | 1 | MT (A9111-AB1) | | |
| | | | 2° pAb donkey anti-rabbit IgG-B1-p2 | 1 | MT (A9230-ZB1) | B1-Alexa546 | 5, S17, S18 |
| | | | 2° pAb goat anti-mouse IgG1-B9-p1 | 1 | MT (A9111-AB9) | | |
| | | | 2° pAb donkey anti-rabbit IgG-B9-p2 | 1 | MT (A9230-ZB9) | B9-Alexa647 | 5, S17, S18 |
| *H. sapiens sapiens* | HeLa cells | $\alpha$-tubulin:$\beta$-tubulin | 1° mAb mouse IgG1 anti-$\beta$-tubulin | 1:1000 | Abc (ab231082) | | |
| | | | 2° pAb goat anti-mouse IgG1-B1-p1 | 1 | MT (A9111-AB1) | | |
| | | | 1° mAb rabbit IgG anti-$\alpha$-tubulin | 1:350 | Abc (ab176560) | | |
| | | | 2° pAb donkey anti-rabbit IgG-B1-p2 | 1 | MT (A9230-ZB1) | B1-Alexa546 | 6, S19, S21 |

**Table S1. Organism, sample type, target protein, 1°Ab probe details, 2°Ab probe details, HCR amplifier details, and figure numbers for HCR imaging of protein:protein complexes.** Split-initiator secondary antibody probes and antibody buffer were obtained from Molecular Technologies (MT) within the Beckman Institute at Caltech. Abc: Abcam. CST: Cell Signaling Technology. Inv: Invitrogen. Sy: Synaptic Systems. Aves: Aves Labs. Pr: Proteintech. Atlas: Atlas Antibodies. AO: antibodies-online.

#### S1.2 Probe and amplifier details for RNA targets using HCR RNA-ISH

| Species | Sample | RNA target | Split-initiator DNA probe pairs | Supplier (catalog #) | HCR amplifier | Figures |
|---|---|---|---|---|---|---|
| *H. sapiens sapiens* | HeLa cells | *U6* | 2 | MT (3882/E032) | B3-Alexa647 | 6, S19, S22 |

**Table S2. Organism, sample type, target RNA, probe set details, HCR amplifier details, and figure numbers for HCR RNA-ISH.** For HCR RNA-ISH, HCR probe sets, amplifiers, and buffers (probe hybridization buffer, probe wash buffer, amplification buffer) were obtained from Molecular Technologies (MT) within the Beckman Institute at Caltech.

#### S1.3 Probe and amplifier details for protein targets using HCR IHC

| Species | Sample | Protein target | 1°Ab probe (unlabeled)<br>2°Ab probe (initiator-labeled) | Dilution factor<br>Working concentration ($\mu$g/mL) | Supplier (catalog #) | HCR amplifier | Figures |
|---|---|---|---|---|---|---|---|
| *H. sapiens sapiens* | HeLa cells | HSP60 | 1°mAb mouse IgG2a anti-HSP60 | 1:50 | Inv (MA3-012) | | |
| | | | 2°pAb goat anti-mouse IgG2a-B5 | 1 | MT (A9112-B5) | B5-Alexa488 | 6, S19, S20 |

**Table S3. Organism, sample type, target protein, 1°Ab probe details, 2°Ab probe details, HCR amplifier details, and figure numbers for HCR IHC.** For HCR IHC, initiator-labeled secondary antibody probes and antibody buffer were obtained from Molecular Technologies (MT) within the Beckman Institute at Caltech. Inv: Invitrogen.

#### S1.4 Oligonucleotide sequences

| HCR System | Oligonucleotide | Length (nt) | Sequence (5' to 3') |
|---|---|---|---|
| B1 | p1 probe | 49 | GAGGAGGGCAGCAAACGGTCGCCCATGTGTACCCGAAATTCAAGTCAGC |
| | p2 probe | 49 | TTCCTGATTCTATTACTTGCCTGATCCCTATGAAGAGTCTTCCTTTACG |
| | proximity probe | 50 | CTTGAATTTCGGGTACACATGGGCTAAGGGATCAGGCAAGTAATAGAATC |
| B6 | p1 probe | 49 | GCAAACTAAACATCCCACAAGGTGAAAGCTGGTACGAATAAGACTACGC |
| | p2 probe | 49 | CTGCACCGGTATATGTTCTGAAGGTGATGCATCCAACTCTAACTAAATC |
| | proximity probe | 50 | GTCTTATTCGTACCAGCTTTCACCACCATCACCTTCAGAACATATACCGG |
| B9 | p1 probe | 49 | CTAACAATCTAAACATACTCGAGGGTGCGGTCTATTCTATTTCCAACGT |
| | p2 probe | 49 | TATTGCGTGTTAGGTGAGTTTGAGATTTGTACACGCCCAAGAACATAAA |
| | proximity probe | 50 | GGAAATAGAATAGACCGCACCCTCACCAAATCTCAAACTCACCTAACACG |

**Table S4. Oligonucleotide sequences for p1, p2, and proximity probe.** Fractional initiators denoted in green (18 nt). Proximity domains denoted in red (24 nt).

##### S1.5 Split-initiator primary antibody probes vs split-initiator secondary antibody probes

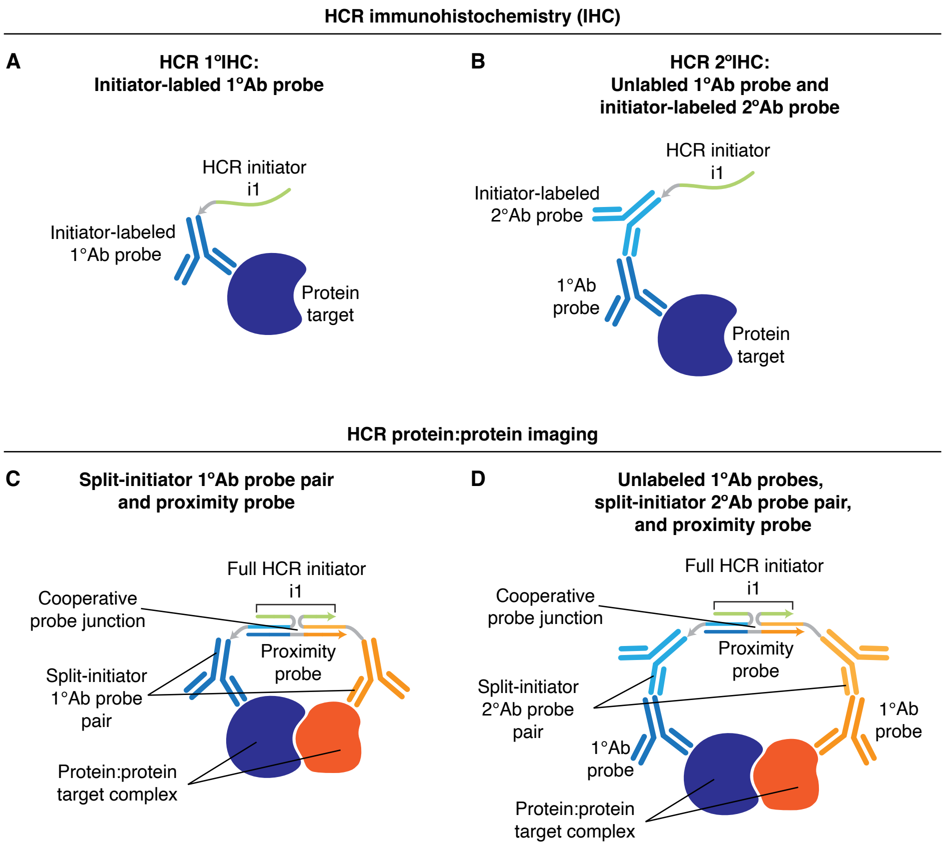

**Figure S1.** Comparison of split-initiator primary antibody probes vs split-initiator secondary antibody probes for imaging protein:protein target complexes. (A,B) HCR immunohistochemistry (Schwarzkopf *et al.*, 2021) using either: HCR 1°IHC with initiator-labeled primary antibody probes (panel A), or HCR 2°IHC using unlabeled primary antibody probes and initiator-labeled secondary antibody probes (panel B). (C,D) HCR imaging of protein:protein complexes using either: a split-initiator primary antibody probe pair and a proximity probe (panel C), or unlabeled primary antibody probes, a split-initiator secondary antibody probe pair, and a proximity probe (panel D).

#### S1.6 Microscope settings

| Sample | Target complex or molecule | Objective | Fluorophore | Laser (nm) | Detector | Detection wavelength (nm) | Pixel size ($x \times y \times z$ $\mu$m) | Figures |
|---|---|---|---|---|---|---|---|---|
| A-431 or HeLa cells | $\beta$-catenin:E-cadherin | 63× | Alexa647 | 638 | HyD X 4 | 648–700 | 0.180 × 0.180 × 0.8 | 3AB, S2, S3 |
| | — | | DAPI | 405 | HyD S 1 | 415–440 | 0.180 × 0.180 × 0.8 | 3AB, S2, S3 |
| Scid.adh.2C2 ProT cells | RUNX1:PU.1 | 63× | Alexa647 | 638 | HyD X 4 | 648–700 | 0.180 × 0.180 × 0.8 | 3CD, S4, S5 |
| | — | | DAPI | 405 | HyD S 1 | 415–440 | 0.180 × 0.180 × 0.8 | 3CD, S4, S5 |
| FFPE human breast tissue | $\beta$-catenin:E-cadherin | 20× | Alexa647 | 638 | HyD X 4 | 648–700 | 0.568 × 0.568 × 3.3 | 3EF, S6, S7 |
| | — | | DAPI | 405 | HyD S 1 | 415–440 | 0.568 × 0.568 × 3.3 | 3EF, S6, S7 |
| A-431 cells | $\beta$-tubulin:$\alpha$-tubulin | 63× | Alexa488 | 488 | HyD S 2 | 495–535 | 0.180 × 0.180 × 0.8 | 4, S8, S9 |
| | $\beta$-catenin:E-cadherin | | Alexa546 | 561 | HyD S 3 | 570–605 | 0.180 × 0.180 × 0.8 | 4, S8, S10 |
| | SC35:SON | | Alexa647 | 638 | HyD X 4 | 648–700 | 0.180 × 0.180 × 0.8 | 4, S8, S11 |
| | — | | DAPI | 405 | HyD S 1 | 415–440 | 0.180 × 0.180 × 0.8 | 4, S8–S11 |
| A-431 cells | $\beta$-catenin:E-cadherin | 63× | Alexa546 | 561 | HyD S 3 | 570–605 | 0.180 × 0.180 × 0.8 | 5, S12, S15 |
| | $\beta$-catenin:E-cadherin | | Alexa647 | 638 | HyD X 4 | 648–700 | 0.180 × 0.180 × 0.8 | 5, S12, S16 |
| | — | | DAPI | 405 | HyD S 1 | 415–440 | 0.180 × 0.180 × 0.8 | 5, S12–S16 |
| FFPE human breast tissue | $\beta$-catenin:E-cadherin | 20× | Alexa546 | 561 | HyD S 3 | 570–605 | 0.568 × 0.568 × 3.3 | 5, S17, S18 |
| | $\beta$-catenin:E-cadherin | | Alexa647 | 638 | HyD X 4 | 648–700 | 0.568 × 0.568 × 3.3 | 5, S17, S18 |
| | — | | DAPI | 405 | HyD S 1 | 415–440 | 0.568 × 0.568 × 3.3 | 5, S17, S18 |
| HeLa cells | HSP60 | 63× | Alexa488 | 488 | HyD S 2 | 495–535 | 0.180 × 0.180 × 0.8 | 6, S19, S20 |
| | $\beta$-tubulin:$\alpha$-tubulin | | Alexa546 | 561 | HyD S 3 | 570–605 | 0.180 × 0.180 × 0.8 | 6, S19, S21 |
| | *U6* | | Alexa647 | 638 | HyD X 4 | 648–700 | 0.180 × 0.180 × 0.8 | 6, S19, S22 |
| | — | | DAPI | 405 | HyD S 1 | 415–440 | 0.180 × 0.180 × 0.8 | 6, S19–S22 |

**Table S5. Microscope settings.** Confocal microscopy was performed with a Leica Stellaris 8 inverted confocal microscope. Objectives were as follows: HC PL APO 20×/0.75 IMM CORR CS2 (catalog # 11506343), HC PL APO 63×/1.40 OIL CS2 (catalog # 11506350), both utilized with oil immersion.

#### S1.7 Image analysis

We build on an image analysis framework developed over a series of publications (Choi *et al.*, 2010; Choi *et al.*, 2014; Choi *et al.*, 2016; Choi *et al.*, 2018; Trivedi *et al.*, 2018; Schwarzkopf *et al.*, 2021). For convenience, here we provide a self-contained description of the details relevant to the present work.

##### S1.7.1 Raw pixel intensities

The total fluorescence within a pixel is a combination of signal and background. Fluorescent background (BACK) arises from up to four sources in each channel:

- autofluorescence (AF): fluorescence inherent to the sample.
- non-specific amplification (NSA): HCR hairpins that bind non-specifically in the sample.
- non-specific detection (NSD): a) For target molecules: probes that bind non-specifically in the sample and subsequently trigger HCR amplification. b) For target complexes: probes that bind non-specifically in the sample and subsequently trigger HCR amplification, or probes that are bound to a target protein not colocalized within a target complex and subsequently trigger HCR amplification.

Fluorescent signal (SIG) in each channel corresponds to:

- signal (SIG): probes that bind specifically in the sample and subsequently trigger HCR amplification via a full HCR initiator.

For pixel $i$ of replicate sample $n$, we denote the background

$$X_{n,i}^{\mathrm{BACK}} = X_{n,i}^{\mathrm{NSD}} + X_{n,i}^{\mathrm{NSA}} + X_{n,i}^{\mathrm{AF}}, \tag{S1}$$

the signal:

$$X_{n,i}^{\mathrm{SIG}}, \tag{S2}$$

and the total fluorescence (SIG+BACK):

$$X_{n,i}^{\mathrm{SIG+BACK}} = X_{n,i}^{\mathrm{SIG}} + X_{n,i}^{\mathrm{BACK}}. \tag{S3}$$

##### S1.7.2 Measurement of signal, background, and signal-to-background for HCR imaging of protein:protein complexes, HCR 2°IF, and HCR RNA-ISH

Background and signal are characterized differently depending on the sample type and protein:protein complex detected:

- For cells on a slide, signal plus background (SIG+BACK) is characterized for pixels in a representative rectangular region of high protein:protein complex expression using an experiment of Type 1a (Table S6) employing the full protocol. For cells for which a suitable control is available in which the protein:protein complex is present at low levels or is absent, background (BACK) can be characterized for pixels in a representative rectangular region of maximum intensity using an experiment of Type 1b (Table S6) employing the full protocol. Alternatively, BACK is characterized for pixels in a representative rectangular region of maximum intensity using experiments of Type 2a and Type 2b (Table S6) in which the interaction probe and 1° Ab against one protein of the protein:protein interaction are omitted, and BACK is estimated as the sum from the two background experiments (BACK $\approx$ $\mathrm{NSD_{p1}}$ + $\mathrm{NSD_{p2}}$ + 2*NSA + 2*AF), which provides an overestimate of background.
- For FFPE human breast tissue sections, signal plus background (SIG+BACK) is characterized for pixels in a representative rectangular region of high protein:protein complex expression using an experiment of Type 1a (Table S6) employing the full protocol, and background (BACK) is characterized for pixels in a representative rectangular region of a different sample with no or low protein:protein complex expression using an experiment of Type 1b (Table S6). Alternatively, SIG+BACK is characterized for pixels in a representative

rectangular region of high protein:protein complex expression using an experiment of Type 3 (Table S6) employing the full protocol, and background (BACK) is characterized for pixels in a representative rectangular region in the same sample with no or low protein:protein complex expression (Table S6).

For HCR IHC experiments, signal plus background (SIG+BACK) is characterized for pixels in a representative rectangular region of high protein expression using an experiment of Type 1 (Table S7) employing the full protocol. Background (BACK) is characterized for pixels in a representative rectangular region of maximum intensity using an experiment of Type 2 (Table S7) in which the $1^\circ$Ab probe is omitted, yielding the partial background estimate BACK $\approx$ $NSD_{2^\circ}$ + NSA + AF.

For HCR RNA-ISH experiments, signal plus background (SIG+BACK) is characterized for pixels in a representative rectangular region of high RNA expression using an experiment of Type 1 (Table S8) employing the full protocol. Background (BACK) is characterized for pixels in a representative rectangular region of maximum intensity using an experiment of Type 2 (Table S8) in which the split-initiator probes are omitted, yielding the partial background estimate BACK $\approx$ NSA + AF.

For the pixels in these regions, we characterize the distribution by plotting an intensity histogram and characterize average performance by calculating the mean pixel intensities ($\bar{X}_n^{\text{BACK}}$ or $\bar{X}_n^{\text{SIG+BACK}}$ for replicate $n$). Performance across replicates is characterized by calculating the sample means ($\bar{X}^{\text{BACK}}$ and $\bar{X}^{\text{SIG+BACK}}$) and standard error of the means ($s_{\bar{X}^{\text{BACK}}}$ and $s_{\bar{X}^{\text{SIG+BACK}}}$). The mean signal is then estimated as

$$\bar{X}^{\text{SIG}} = \bar{X}^{\text{SIG+BACK}} - \bar{X}^{\text{BACK}} \tag{S4}$$

with the standard error of the mean estimated via uncertainty propagation as

$$s_{\bar{X}^{\text{SIG}}} \leq \sqrt{(s_{\bar{X}^{\text{SIG+BACK}}})^2 + (s_{\bar{X}^{\text{BACK}}})^2}. \tag{S5}$$

The upper bound on estimated standard error of the mean holds under the assumption that the correlation between SIG+BACK and BACK is non-negative. The mean signal-to-background ratio is estimated as:

$$\bar{X}^{\text{SIG/BACK}} = \bar{X}^{\text{SIG}} / \bar{X}^{\text{BACK}} \tag{S6}$$

with standard error of the mean estimated via uncertainty propagation as

$$s^{\text{SIG/BACK}} \leq \bar{X}^{\text{SIG/BACK}} \sqrt{\left(\frac{s_{\bar{X}^{\text{SIG}}}}{\bar{X}^{\text{SIG}}}\right)^2 + \left(\frac{s_{\bar{X}^{\text{BACK}}}}{\bar{X}^{\text{BACK}}}\right)^2}. \tag{S7}$$

The upper bound on estimated standard error of the mean holds under the assumption that the correlation between SIG and BACK is non-negative.

| Experiment type | Quantity | Reagents | | | | | | Complex |
|---|---|---|---|---|---|---|---|---|
| | | 1° Ab for protein 1 | 1° Ab for protein 2 | 2° Ab probe p1 | 2° Ab probe p2 | Proximity probe | Amplifier | expression region in sample |
| 1a | SIG+NSD+NSA+AF = SIG+BACK | ✓ | ✓ | ✓ | ✓ | ✓ | ✓ | high |
| 1b | NSD+NSA+AF = BACK | ✓ | ✓ | ✓ | ✓ | ✓ | ✓ | no/low |
| 2a | $NSD_{P1}$+NSA+AF | ✓ | | ✓ | | ✓ | ✓ | high |
| 2b | $NSD_{P2}$+NSA+AF | | ✓ | | ✓ | ✓ | ✓ | high |
| 3 | SIG+NSD+NSA+AF = SIG+BACK | ✓ | ✓ | ✓ | ✓ | ✓ | ✓ | high |
| 3 | NSD+NSA+AF = BACK | ✓ | ✓ | ✓ | ✓ | ✓ | ✓ | no/low |

**Table S6. Experiment types for HCR protein:protein imaging using unlabeled primary antibody probes, split-initiator secondary antibody probes, and a proximity probe.** Characterize signal, background, and signal-to-background.

| Experiment type | Quantity | Reagents | | | Expression region |
|---|---|---|---|---|---|
| | | 1° Ab | 2° Ab-initiator | Amplifier | in sample |
| 1 | SIG+NSD+NSA+AF = SIG+BACK | ✓ | ✓ | ✓ | high |
| 2 | $NSD_{2^\circ}$+NSA+AF | | ✓ | ✓ | high |

**Table S7. Experiment types for HCR IHC using unlabeled primary antibody probes and initiator-labeled secondary antibody probes.** Characterize signal, background, and signal-to-background.

| Experiment type | Quantity | Reagents | | Expression region |
|---|---|---|---|---|
| | | Split-initiator probes | Amplifier | in sample |
| 1 | SIG+NSD+NSA+AF = SIG+BACK | ✓ | ✓ | high |
| 2 | NSA+AF | | ✓ | high |

**Table S8. Experiment types for HCR RNA-ISH using split-initiator DNA probes.** Characterize signal, background, and signal-to-background.

##### S1.7.3 Normalized voxel intensities for qHCR imaging: protein:protein complex relative quantitation with subcellular resolution in an anatomical context

For quantitative imaging using HCR, precision increases with voxel size as long as the imaging voxels remain smaller than the features in the expression pattern (see Section S2.2 of (Trivedi *et al.*, 2018)). To increase precision, we calculate raw voxel intensities by averaging neighboring pixel intensities while still maintaining a subcellcular voxel size. To facilitate relative quantitation between voxels, we estimate the normalized HCR signal of voxel $j$ in replicate $n$ as:

$$x_{n,j} \equiv \frac{X_{n,j}^{\mathrm{SIG+BACK}} - X^{\mathrm{BOT}}}{X^{\mathrm{TOP}} - X^{\mathrm{BOT}}}, \tag{S8}$$

which translates and rescales the data so that the voxel intensities in each channel fall in the interval [0,1]. Here,

$$X^{\mathrm{BOT}} \equiv \bar{X}^{\mathrm{BACK}} \tag{S9}$$

is the mean background across replicates (see Section S1.7.2) and

$$X^{\mathrm{TOP}} \equiv \max_{n,j} X_{n,j}^{\mathrm{SIG+BACK}} \tag{S10}$$

is the maximum total fluorescence for a voxel across replicates.

Pairwise expression scatter plots that each display normalized voxel intensities for two channels (e.g., Figures 4 and 5 of (Trivedi *et al.*, 2018)) provide a powerful quantitative framework for performing multidimensional read-out/read-in analyses (Figure 6 of (Trivedi *et al.*, 2018)). Read-out from anatomical space to expression space enables discovery of expression clusters of voxels with quantitatively related expression levels and ratios (amplitudes and slopes in the expression scatter plots), while read-in from expression space to anatomical space enables discovery of the corresponding anatomical locations of these expression clusters within the sample. The simple and practical normalization approach of (S8)–(S10) translates and rescales all voxels identically within a given channel (enabling comparison of amplitudes and slopes in scatter plots between replicates), and does not attempt to remove scatter in the normalized signal estimate that is caused by scatter in background.

To validate qHCR protein:protein complex imaging with subcellular resolution in human cells ($2\times2\times0.8$ $\mu$m voxels) and FFPE human breast tissue sections ($2\times2\times3.3$ $\mu$m voxels), Figures 5C, S12C, and S17C display highly correlated normalized voxel intensities for 2-channel redundant detection of a protein:protein complex. In this setting, accuracy corresponds to linearity with zero intercept, and precision corresponds to scatter around the line (Trivedi *et al.*, 2018).

#### S1.8 Protocols for protein:protein complex imaging in human cells

##### S1.8.1 Preparation of fixed adherent human cells on a chambered slide

1. Coat the bottom of each chamber by applying 100 $\mu$L of 0.01% poly-D-lysine prepared in molecular biology grade $H_2O$.
   NOTE: *For each step, a volume of 100 $\mu$L is sufficient per chamber on an 18-chamber slide. Scale volumes accordingly if using a different chambered format.*

2. Incubate for at least 30 min at room temperature.

3. Aspirate the coating solution and wash each chamber 2× with molecular biology grade $H_2O$.

4. Add the desired number of cells to each chamber.

5. Grow the cells to the desired confluency for 24–48 h.

6. Aspirate the growth media and wash each chamber with DPBS.
   NOTE: *Avoid using DPBS with calcium chloride and magnesium chloride, as this leads to increased autofluorescence.*

7. Add 4% formaldehyde in PBS to each chamber.
   CAUTION: *Use formaldehyde with extreme care, as it is a hazardous material.*

8. Incubate for 10 min at room temperature.

9. Remove fixative and wash each chamber 2× with DPBS.

10. Aspirate DPBS and add ice-cold 70% ethanol (EtOH) to permeabilize the cells.

11. Permeabilize cells for 3 h at 4 °C.
    NOTE: *Alternatively, the cells may be permeabilized overnight (or longer) at -20 °C.*

##### S1.8.2 HCR protein:protein complex imaging with/without HCR IHC and HCR RNA-ISH using simultaneous HCR signal amplification for all targets

**Protein detection stage**

1. Aspirate EtOH from sample and wash 2 × 5 min with 1× PBS.
2. Add antibody buffer to the sample. Incubate for 1 h at room temperature with gentle agitation.
3. Prepare working concentration of primary antibodies in antibody buffer.
   NOTE: *Follow the manufacturer's guidelines for the primary antibody working concentration.*
4. Replace antibody buffer with primary antibody solution and incubate overnight (>12 h) at 4 °C with gentle agitation.
   NOTE: *Incubation may be optimized (e.g., 1–2 h at room temperature) depending on the antibody used.*
5. Remove excess antibodies by washing 3 × 5 min with 1× PBST at room temperature with gentle agitation.
6. Prepare secondary antibody probe solution containing split-initiator secondary antibody probes (p1 and p2), and optionally initiator-labeled secondary antibodies for HCR IHC, at 1 $\mu$g/mL in antibody buffer.
7. Add the secondary antibody probe solution to the sample. Incubate for 1 h at room temperature with gentle agitation.
8. Remove excess secondary antibody probes by washing 3 × 5 min with 1× PBST at room temperature with gentle agitation.
9. Wash with 5× SSCT for 5 min at room temperature.
10. Add amplification buffer. Incubate for 30 min at room temperature.
    NOTE: *Equilibrate amplification buffer to room temperature before use.*
11. Prepare a 1.6 nM proximity probe solution in amplification buffer at room temperature.
12. Remove amplification buffer and add proximity probe solution. Incubate for 1 h at room temperature.
13. Wash 3 × 5 min with 5× SSCT at room temperature.
14. Proceed to **RNA detection stage** for co-detection of RNA. Otherwise, proceed to **Amplification stage**.

**RNA detection stage**

1. Post-fix sample with 4% formaldehyde in PBS. Incubate for 10 min at room temperature.
   CAUTION: *Use formaldehyde with extreme care, as it is a hazardous material.*
2. Wash 3 × 5 min with 1× PBST at room temperature.
3. Wash with 5× SSCT for 5 min at room temperature.
4. Add probe hybridization buffer. Incubate for 30 min at 37 °C.
   CAUTION: *Probe hybridization buffer contains formamide, a hazardous material.*
   NOTE: *Pre-heat probe hybridization buffer to 37 °C before use.*
5. Prepare a 16 nM probe solution in probe hybridization buffer at 37 °C.
6. Remove the probe hybridization buffer and add the probe solution.
7. Incubate overnight (>12 h) at 37 °C.

8. Remove excess probes by washing 4 × 5 min with probe wash buffer at 37 °C.
   CAUTION: *Probe wash buffer contains formamide, a hazardous material.*
   NOTE: *Pre-heat probe wash buffer to 37 °C before use.*

9. Wash 3 × 5 min with 5× SSCT at room temperature.

10. Proceed to **Amplification stage**.

**Amplification stage**

1. Add amplification buffer. Incubate for 30 min at room temperature.
   NOTE: *Equilibrate amplification buffer to room temperature before use.*

2. Separately prepare hairpin h1 and hairpin h2 by snap cooling (heat at 95 °C for 90 seconds and cool to room temperature in a dark drawer for 30 min).
   NOTE: *HCR hairpins h1 and h2 are provided in hairpin storage buffer ready for snap cooling. h1 and h2 should be snap cooled in separate tubes.*

3. Prepare a 60 nM hairpin solution by adding snap-cooled h1 hairpins and snap-cooled h2 hairpins to amplification buffer at room temperature.

4. Remove the amplification buffer and add the hairpin solution.

5. Incubate overnight (>12 h) protected from light at room temperature.

6. Remove excess hairpins by washing 5 × 5 min with 5× SSCT at room temperature.

**Sample mounting for microscopy**

1. Add DAPI Fluoromount-G mounting medium.

2. The coverslip can be stored at 4 °C protected from light prior to imaging.
   NOTE: *see Section S1.6 for details of confocal microscopy used to image human cells on a chambered coverslip.*

##### S1.8.3 Buffers for HCR protein:protein complex imaging with/without HCR IHC and HCR RNA-ISH

HCR probes (split-initiator antibody probes, initiator-labeled antibody probes, split-initiator DNA probes), amplifiers, and buffers (antibody buffer, probe hybridization buffer, probe wash buffer, amplification buffer) are available from Molecular Technologies. Probe hybridization buffer and probe wash buffer should be stored at -20 °C. Antibody buffer and amplification buffer should be stored at 4 °C. Make sure all solutions are well mixed before use.

| **PBST** | For 500 mL of solution |
|---|---|
| 1× PBS | 50 mL of 10× PBS |
| 0.1% Tween-20 | 5 mL of 10% Tween-20 |
| | Fill up to 500 mL with ultrapure $H_2O$ |
| | Filter with a 0.2 $\mu$m Nalgene Rapid-Flow filter |

| **5× SSCT** | For 500 mL of solution |
|---|---|
| 5× saline sodium citrate (SSC) | 125 mL of 20× SSC |
| 0.1% Tween-20 | 5 mL of 10% Tween-20 |
| | Fill up to 500 mL with ultrapure $H_2O$ |
| | Filter with a 0.2 $\mu$m Nalgene Rapid-Flow filter |

##### S1.8.4 Reagents and supplies

ibidi $\mu$-slide 18 well ibiTreat (ibidi, 81816)
Poly-D-lysine hydrobromide (Sigma-Aldrich, P7280)
Molecular biology grade $H_2O$ (Corning, 46-000-CV)
DPBS, no calcium, no magnesium (Gibco, 14190144)
Image-iT 4% formaldehyde fixative solution in PBS (methanol-free) (Invitrogen, FB002)
100% EtOH (Koptec, V1001)
10× PBS (Invitrogen, AM9624)
20× Saline sodium citrate (SSC) (Invitrogen, 15557-044)
10% Tween-20 (Teknova, T0710)
DAPI Fluoromount-G (SouthernBiotech, 0100-20)

#### S1.9 Protocols for protein:protein complex imaging in mouse proT cells

To revert Scid.adh.2C2 (Dionne *et al.*, 2005) cells to an earlier pre-commitment T cell stage, the PU.1 protein is exogenously introduce by retroviral transduction as previously described (Hosokawa *et al.*, 2018). As a control, Scid.adh.2C2 cells are also retrovirally transduced with a control vector that does not encode for the PU.1 protein.

##### S1.9.1 Retroviral packaging of vector DNA

1. Culture Phoenix-ECO cells in Dulbecco's Modified Eagle Medium (DMEM) supplemented with 10% fetal bovine serum (FBS) and 1× Penicillin-Streptomycin-Glutamine.
2. To perform retroviral vector DNA packaging, transfect Phoenix-ECO cells with retroviral vector DNA encoding for the PU.1 protein (or, as a control, with retroviral vector DNA not encoding for the PU.1 protein) via the FuGENE 6 transfection reagent according to the manufacturer's instructions.
3. After 48–72 h, collect the cell supernatant, which contains the packaged retrovirus, and filter through a 0.45 $\mu$m syringe filter.
4. Store the filtered retrovirus supernatant in 1 mL aliquots at -80 °C.

##### S1.9.2 Retroviral infection of mouse proT cells

1. Coat a non-treated 24-well plate with 300–500 $\mu$L of 50 $\mu$g/mL RetroNectin overnight at 4 °C.
2. Remove excess RetroNectin from the 24-well plate.
3. Wash each well with 500 $\mu$L 1× phosphate-buffered saline (PBS).
4. Thaw retrovirus supernatant in a 37 °C water bath.
5. Remove 1× PBS from the 24-well plate and add 1 mL retrovirus supernatant to each well.
6. Centrifuge the plate at 2000 rcf for 2 h in a pre-heated 32 °C centrifuge.
7. Aspirate the liquid from each well.
8. Add $1\times10^5$ Scid.adh.2C2 cells in 1 mL culture media to each well.
9. Centrifuge the plate at 250 rcf for 5 min in a pre-heated 32 °C centrifuge.
10. Incubate overnight at 37 °C in a 5% $CO_2$ incubator.
11. The next day, scrape Scid.adh.2C2 cells from the surface of the 24-well plate.
12. Add Scid.adh.2C2 cells at a concentration of $1\times10^5$/mL to a new tissue culture-treated 24-well plate.
13. Incubate overnight at 37 °C in a 5% $CO_2$ incubator.
14. Remove Scid.adh.2C2 cells from the surface of the 24-well plate by pipetting.
15. Centrifuge Scid.adh.2C2 cells at 250 rcf for 7 min in a pre-cooled 4°C centrifuge.
16. Remove supernatant from the Scid.adh.2C2 cells and resuspend the cells in FACS buffer to a concentration of $1\times10^4$ cells/$\mu$L.
17. Add biotinylated anti-hNGFR (human NGFR) antibody to the cell solution to reach a final antibody dilution factor of 1:300. Mix gently.

18. Incubate the cell solution for 30 min on ice.

19. Centrifuge Scid.adh.2C2 cells at 250 rcf for 7 min in a pre-cooled 4°C centrifuge. Remove the supernatant.

20. Wash the cells twice by adding 300 $\mu$L FACS buffer, centrifuging at 250 rcf for 7 min in a pre-cooled 4°C centrifuge, and removing the supernatant.

21. Label hNGFR+ cells with streptavidin microbeads according to the manufacturer's instructions.

22. Enrich for hNGFR-positive cells by using MACS LS magnetic columns according to the manufacturer's instructions.

23. Centrifuge Scid.adh.2C2 cells at 250 rcf for 7 min in a pre-cooled 4°C centrifuge.

24. Remove supernatant from the Scid.adh.2C2 cells and resuspend the cells in FACS buffer to a concentration of $1\times10^4$ cells/$\mu$L.

25. Add Alexa488-labeled anti-CD45 antibody to the cell solution to reach a final antibody dilution factor of 1:600. Mix gently.

26. Incubate the cell solution for 30 min on ice protected from light.

27. Centrifuge Scid.adh.2C2 cells at 250 rcf for 7 min in a pre-cooled 4°C centrifuge. Remove the supernatant.

28. Wash the cells twice by adding 300 $\mu$L FACS buffer, centrifuging at 250 rcf for 7 min in a pre-cooled 4°C centrifuge, and removing the supernatant.

29. Resuspend the cells in 1× PBS (Gibco) and bring to a concentration of ~2,777 cells/$\mu$L.

##### S1.9.3 Preparation of fixed mouse proT cells on a chambered slide

1. Prepare a No. 1.5 coverslip by washing with 100% ethanol (EtOH) and drying.

2. Coat the coverslip by applying 1.5 mL of 0.01% poly-D-lysine for 1 h.

3. Aspirate the coating solution and wash the coverslip 3× with 1× PBS (Gibco). Air dry the coverslip after the final wash.

4. Affix a microchamber flow cell to the coverslip.

5. Add 18 $\mu$L of the ~2,777 cells/$\mu$L solution to each microchamber port (~50,000 cells/port).
   NOTE: *For each step, a volume of 18 $\mu$L is sufficient for each microchamber of the flow cell. Scale volumes accordingly if using a different chambered format.*

6. Centrifuge the coverslip at 250 rcf for 5 min in a pre-cooled 4°C centrifuge to adhere the cells to the coverslip.

7. Aspirate the 1× PBS.

8. Add 4% formaldehyde in PBS to each chamber.
   CAUTION: *Use formaldehyde with extreme care, as it is a hazardous material.*

9. Incubate for 10 min at room temperature.

10. Remove fixative and wash each chamber 2× with DPBS.

11. Aspirate DPBS and add ice-cold 70% ethanol (EtOH) to permeabilize the cells.

12. Permeabilize cells for 3 h at -20 °C in a humidified chamber.
    NOTE: *Keep the coverslip in a humidified chamber for all future steps to prevent evaporation.*

13. Transfer cells to 4 °C and further permeabilize for 20 min.

##### S1.9.4 HCR protein:protein complex imaging

###### Protein detection stage

1. Aspirate EtOH from sample and wash 2 × 5 min with 1× PBS.
2. Add antibody buffer to the sample. Incubate for 1 h at room temperature with gentle agitation.
3. Prepare working concentration of primary antibodies in antibody buffer.
   NOTE: *Follow the manufacturer's guidelines for the primary antibody working concentration.*
4. Replace antibody buffer with primary antibody solution and incubate overnight (>12 h) at 4 °C with gentle agitation.
   NOTE: *Incubation may be optimized (e.g., 1–2 h at room temperature) depending on the antibody used.*
5. Remove excess antibodies by washing 3 × 5 min with 1× PBST at room temperature with gentle agitation.
6. Prepare secondary antibody probe solution containing split-initiator secondary antibody probes (p1 and p2) at 1 $\mu$g/mL in antibody buffer.
7. Add secondary antibody probe solution to the sample. Incubate for 1 h at room temperature with gentle agitation.
8. Remove excess secondary antibody probes by washing 3 × 5 min with 1× PBST at room temperature with gentle agitation.
9. Wash with 5× SSCT for 5 min at room temperature.
10. Add amplification buffer. Incubate for 30 min at room temperature.
    NOTE: *Equilibrate amplification buffer to room temperature before use.*
11. Prepare a 1.6 nM proximity probe solution in amplification buffer at room temperature.
12. Remove amplification buffer and add proximity probe solution. Incubate for 1 h at room temperature.
13. Wash 3 × 5 min with 5× SSCT at room temperature.

###### Amplification stage

1. Add amplification buffer. Incubate for 30 min at room temperature.
   NOTE: *Equilibrate amplification buffer to room temperature before use.*
2. Separately prepare hairpin h1 and hairpin h2 by snap cooling (heat at 95 °C for 90 seconds and cool to room temperature in a dark drawer for 30 min).
   NOTE: *HCR hairpins h1 and h2 are provided in hairpin storage buffer ready for snap cooling. h1 and h2 should be snap cooled in separate tubes.*
3. Prepare a 60 nM hairpin solution by adding snap-cooled h1 hairpins and snap-cooled h2 hairpins to amplification buffer at room temperature.
4. Remove the amplification buffer and add the hairpin solution.
5. Incubate overnight (>12 h) protected from light at room temperature.
6. Remove excess hairpins by washing 5 × 5 min with 5× SSCT at room temperature.

**Sample mounting for microscopy**

1. Add DAPI Fluoromount-G mounting medium.
2. The coverslip can be stored at 4 °C protected from light prior to imaging.
   NOTE: *see Section S1.6 for details of confocal microscopy used to image mouse proT cells on a chambered slide.*

##### S1.9.5 Buffers for HCR protein:protein complex imaging in mouse proT cells

HCR probes, amplifiers, and buffers (antibody buffer and amplification buffer) are available from Molecular Technologies. Antibody buffer and amplification buffer should be stored at 4 °C. Make sure all solutions are well mixed before use.

| **FACS buffer** | For 50 mL of solution |
|---|---|
| 1× Hanks' balanced salt solution (HBSS) | 49.5 mL of 1× HBSS |
| 10 mM HEPES | 0.5 mL of 1 M HEPES |
| 0.5% bovine serum albumin (BSA) | 250 mg of BSA |

| **PBST** | For 500 mL of solution |
|---|---|
| 1× PBS | 50 mL of 10× PBS (Invitrogen) |
| 0.1% Tween-20 | 5 mL of 10% Tween-20 |
| | Fill up to 500 mL with ultrapure $H_2O$ |
| | Filter with a 0.2 $\mu$m Nalgene Rapid-Flow filter |

| **5× SSCT** | For 500 mL of solution |
|---|---|
| 5× saline sodium citrate (SSC) | 125 mL of 20× SSC |
| 0.1% Tween-20 | 5 mL of 10% Tween-20 |
| | Fill up to 500 mL with ultrapure $H_2O$ |
| | Filter with a 0.2 $\mu$m Nalgene Rapid-Flow filter |

##### S1.9.6 Reagents and supplies

Phoenix-ECO cells (ATCC, CRL-3214)
DMEM (Gibco, 12430-054)
Fetal bovine serum (FBS) (Sigma-Aldrich, F2442)
Penicillin-Streptomycin-Glutamine (100X) (Gibco, 10378-016)
FuGENE 6 Transfection Reagent (Promega, E2691)
0.45 $\mu$m syringe filter (Pall, 4614)
Non-treated 24-well plate (Corning, 351147)
RetroNectin (Takara Bio, T100B)
1× PBS (Gibco, 10010-023)
Tissue culture-treated 24-well plate (Corning, 3524)
100% EtOH (Koptec, V1001)
RPMI1640 medium (Gibco, 31800022)
1× HBSS, no calcium, no magnesium, no phenol red (Gibco, 14175-095)
Bovine serum albumin (BSA) (Roche, 03117332001)
HEPES (1 M) (Gibco, 15630-080)
Biotinylated anti-hNGFR antibody (mouse IgG1 mAb) (BioLegend, 345122)
Streptavidin MicroBeads (Miltenyi Biotec, 130-048-101)
MACS LS magnetic columns (Miltenyi Biotec, 130-042-401)
Alexa488-labeled anti-CD45 antibody (rat IgG2b mAb) (BioLegend, 103122)
Poly-D-lysine (Gibco, A38904-01)
Microchambers (Grace Bio-Labs, custom SecureSeal Flowcell, 15 mm x 75 mm OD, 14 x (3 mm x 11 mm) ID, 0.5 mm thick, A2, 0.020" cover with ports)
DPBS, no calcium, no magnesium (Gibco, 14190144)
Image-iT 4% formaldehyde fixative solution in PBS (methanol-free) (Invitrogen, FB002)
10× PBS (Invitrogen, AM9624)

20× Saline sodium citrate (SSC) (Invitrogen, 15557-044)
10% Tween-20 (Teknova, T0710)
DAPI Fluoromount-G (SouthernBiotech, 0100-20)

#### S1.10 Protocols for FFPE human breast tissue sections

##### S1.10.1 Preparation of formalin-fixed paraffin-embedded (FFPE) human breast tissue sections

1. Bake slides in a dry oven for 1 h at 60 °C to improve sample adhesion to the slide.
2. In a fume hood, deparaffinize FFPE tissue by immersing the slide in Pro-Par Clearant for 2 × 5 min. Gently move the slide up and down every minute.
   NOTE: *If desired, a larger number of slides can be processed together using a Coplin jar.*
3. Immerse the slide in 100% ethanol (EtOH) for 2 × 3 min at room temperature. Gently move the slide up and down every minute.
4. Immerse the slide in 95% EtOH for 3 min at room temperature.
5. Immerse the slide in 70% EtOH for 3 min at room temperature.
6. Immerse the slide in nanopure $H_2O$ for 3 min at room temperature.
7. Heat antigen retrieval buffer in a heatproof container with a digital steamer until >98 °C.
8. Immerse the slide in the heated antigen retrieval buffer in the digital steamer for 15 min.
9. Remove the slide from the antigen retrieval buffer and immediately immerse in nanopure $H_2O$ for 10 min at room temperature.
10. Remove the slide and gently tap off excess nanopure $H_2O$.
11. Carefully dry around the tissue with a Kimwipe.
12. Draw a hydrophobic barrier around the tissue with a hydrophobic pen.
13. Place the slide in a humidified chamber.
    NOTE: *Keep the slide in a humidified chamber for all future steps to prevent evaporation.*

##### S1.10.2 HCR protein:protein complex imaging in FFPE human breast tissue sections

###### Protein detection stage

1. Apply antibody buffer to the tissue section. Incubate for 1 h at room temperature.
   NOTE: *Scale volumes according to the size of the tissue. A volume of 100 $\mu$L was utilized here for each step.*

2. Prepare working concentration of primary antibodies in antibody buffer.
   NOTE: *Follow the manufacturer's guidelines for the primary antibody working concentration.*

3. Replace antibody buffer with primary antibody solution and incubate overnight ($>$12 h) at 4 °C.
   NOTE: *Incubation may be optimized (e.g., 1–2 h at room temperature) depending on the antibody used.*

4. Remove excess antibodies by washing 3 $\times$ 5 min with 1$\times$ PBST at room temperature.

5. Prepare secondary antibody probe solution containing split-initiator secondary antibody probes (p1 and p2) at 1 $\mu$g/mL in antibody buffer.

6. Add secondary antibody probe solution to the sample. Incubate for 1 h at room temperature.

7. Remove excess secondary antibody probes by washing 3 $\times$ 5 min with 1$\times$ PBST at room temperature.

8. Wash with 5$\times$ SSCT for 5 min at room temperature.

9. Add amplification buffer. Incubate for 30 min at room temperature.
   NOTE: *Equilibrate amplification buffer to room temperature before use.*

10. Prepare a 1.6 nM proximity probe solution in amplification buffer at room temperature.

11. Remove amplification buffer and add proximity probe solution. Incubate for 1 h at room temperature.

12. Wash 3 $\times$ 5 min with 5$\times$ SSCT at room temperature.

###### Amplification stage

1. Add amplification buffer. Incubate for 30 min at room temperature.
   NOTE: *Equilibrate amplification buffer to room temperature before use.*

2. Separately prepare hairpin h1 and hairpin h2 by snap cooling (heat at 95 °C for 90 seconds and cool to room temperature in a dark drawer for 30 min).
   NOTE: *HCR hairpins h1 and h2 are provided in hairpin storage buffer ready for snap cooling. h1 and h2 should be snap cooled in separate tubes.*

3. Prepare a 60 nM hairpin solution by adding snap-cooled h1 hairpins and snap-cooled h2 hairpins to amplification buffer at room temperature.

4. Remove the amplification buffer and add the hairpin solution.

5. Incubate overnight ($>$12 h) protected from light at room temperature.

6. Remove excess hairpins by washing with 5$\times$ SSCT at room temperature:

   (a) 2 $\times$ 5 min
   (b) 2 $\times$ 15 min
   (c) 1 $\times$ 5 min

**Sample mounting for microscopy**

1. Carefully dry around the section with a Kimwipe.
2. Apply 60 $\mu$L of DAPI Fluoromount-G to the section.
3. Carefully lower a 22 mm $\times$ 30 mm No. 1.5 coverslip on top of the section.
4. Slides can be stored at 4 °C protected from light prior to imaging.
   NOTE: *see Section S1.6 for details of confocal microscopy used to image FFPE human breast tissue sections.*

##### S1.10.3 Buffers for HCR protein:protein complex imaging in FFPE human breast tissue sections

HCR probes, amplifiers, and buffers (antibody buffer and amplification buffer) are available from Molecular Technologies. Antibody buffer and amplification buffer should be stored at 4 °C. Make sure all solutions are well mixed before use.

| **PBST** | For 500 mL of solution |
|---|---|
| 1× PBS | 50 mL of 10× PBS |
| 0.1% Tween-20 | 5 mL of 10% Tween-20 |
| | Fill up to 500 mL with ultrapure $H_2O$ |
| | Filter with a 0.2 $\mu$m Nalgene Rapid-Flow filter |

| **5× SSCT** | For 500 mL of solution |
|---|---|
| 5× saline sodium citrate (SSC) | 125 mL of 20× SSC |
| 0.1% Tween-20 | 5 mL of 10% Tween-20 |
| | Fill up to 500 mL with ultrapure $H_2O$ |
| | Filter with a 0.2 $\mu$m Nalgene Rapid-Flow filter |

##### S1.10.4 Reagents and supplies

Pro-Par Clearant (ANATECH LTD, 510)
100% EtOH (Koptec, V1001)
Antigen retrieval buffer (100X Citrate Buffer) (Abcam, ab93678)
10× PBS (Invitrogen, AM9624)
20× Saline sodium citrate (SSC) (Invitrogen, 15557-044)
10% Tween-20 (Teknova, T0710)
DAPI Fluoromount-G (SouthernBiotech, 0100-20)

### S2 Additional studies

#### S2.1 Summary of signal-to-background estimates for HCR protein:protein imaging, HCR IHC, and HCR RNA-ISH

| Method | Sample | Target | Type | Probes | Amplifier | SIG/BACK | Plex | Figures | Table |
|---|---|---|---|---|---|---|---|---|---|
| HCR protein:protein | human cells on a slide | $\beta$-catenin:E-cadherin | protein:protein | two 1°mAb + two 2°pAb-split-init | B1-Alexa647 | $26 \pm 4$ | 1 | 3A, S2, S3 | S10 |
| HCR protein:protein | mouse proT cells on a slide | RUNX1:PU.1 | protein:protein | one 1°pAb + one 1°mAb + two 2°pAb-split-init | B1-Alexa647 | $15 \pm 3$ | 1 | 3B, S4, S5 | S11 |
| HCR protein:protein | FFPE human breast tissue section | $\beta$-catenin:E-cadherin | protein:protein | two 1°mAb + two 2°pAb-split-init | B1-Alexa647 | $30 \pm 3$ | 1 | 3C, S6, S7 | S12 |
| HCR protein:protein | human cells on a slide | $\beta$-tubulin:$\alpha$-tubulin | protein:protein | one 1°mAb + one 1°pAb + two 2°pAb-split-init | B6-Alexa488 | $32 \pm 4$ | 3 | 4, S8, S9 | S13 |
| HCR protein:protein | human cells on a slide | $\beta$-catenin:E-cadherin | protein:protein | one 1°pAb + one 1°mAb + two 2°pAb-split-init | B1-Alexa546 | $22 \pm 4$ | 3 | 4, S8, S10 | S13 |
| HCR protein:protein | human cells on a slide | SC35:SON | protein:protein | one 1°mAb + one 1°pAb + two 2°pAb-split-init | B9-Alexa647 | $90 \pm 20$ | 3 | 4, S8, S11 | S13 |
| HCR protein:protein | human cells on a slide | $\beta$-catenin:E-cadherin | protein:protein | two 1°mAb + two 2°pAb-split-init | B1-Alexa546 | $16 \pm 2$ | 2 | 5, S12, S15 | S15 |
| HCR protein:protein | human cells on a slide | $\beta$-catenin:E-cadherin | protein:protein | two 1°mAb + two 2°pAb-split-init | B6-Alexa647 | $15 \pm 3$ | 2 | 5, S12, S16 | S15 |
| HCR protein:protein | FFPE human breast tissue section | $\beta$-catenin:E-cadherin | protein:protein | two 1°mAb + two 2°pAb-split-init | B1-Alexa546 | $52 \pm 6$ | 2 | 5, S17, S18 | S17 |
| HCR protein:protein | FFPE human breast tissue section | $\beta$-catenin:E-cadherin | protein:protein | two 1°mAb + two 2°pAb-split-init | B9-Alexa647 | $91 \pm 9$ | 2 | 5, S17, S18 | S17 |
| HCR protein:protein | human cells on a slide | $\alpha$-tubulin:$\beta$-tubulin | protein:protein | two 1°mAb + two 2°pAb-split-init | B1-Alexa546 | $110 \pm 30$ | 3 | 6, S19, S21 | S18 |
| HCR IHC | human cells on a slide | HSP60 | protein | 1°mAb + 2°pAb-init | B5-Alexa488 | $800 \pm 100$ | 3 | 6, S19, S20 | S18 |
| HCR RNA-ISH | human cells on a slide | *U6* | RNA | two DNA-split-init pairs | B3-Alexa647 | $160 \pm 30$ | 3 | 6, S19, S22 | S18 |

**Table S9. Signal-to-background summary for HCR protein:protein imaging with or without HCR IHC and HCR RNA-ISH.** Mean ± SEM for representative regions within $N = 3$ wells on a slide (mammalian cells on a slide) or $N = 3$ replicate sections (FFPE human breast tissue sections).

#### S2.2 Replicates, signal, and background for protein:protein complex imaging in human cells (cf. Figure 3AB)

For the $\beta$-catenin:E-cadherin target complex, A-431 cells serve as the positive sample and provide an estimate of signal + background (SIG+BACK), while HeLa cells serve as the negative sample and provide an estimate of background (BACK). Additional studies are presented as follows:

- Figure S2 displays images for $N = 3$ replicate wells on a multi-well slide for A-431 cells (panel A) and HeLa cells (panel B) (cf. Figure 3AB).
- Figure S3 displays representative regions used for measurement of signal and background.
- Table S10 displays estimated values for signal, background, and signal-to-background.

**Protocol:** Section S1.8.
**Sample:** A-431 cells and HeLa cells.
**Reagents:** Table S1.

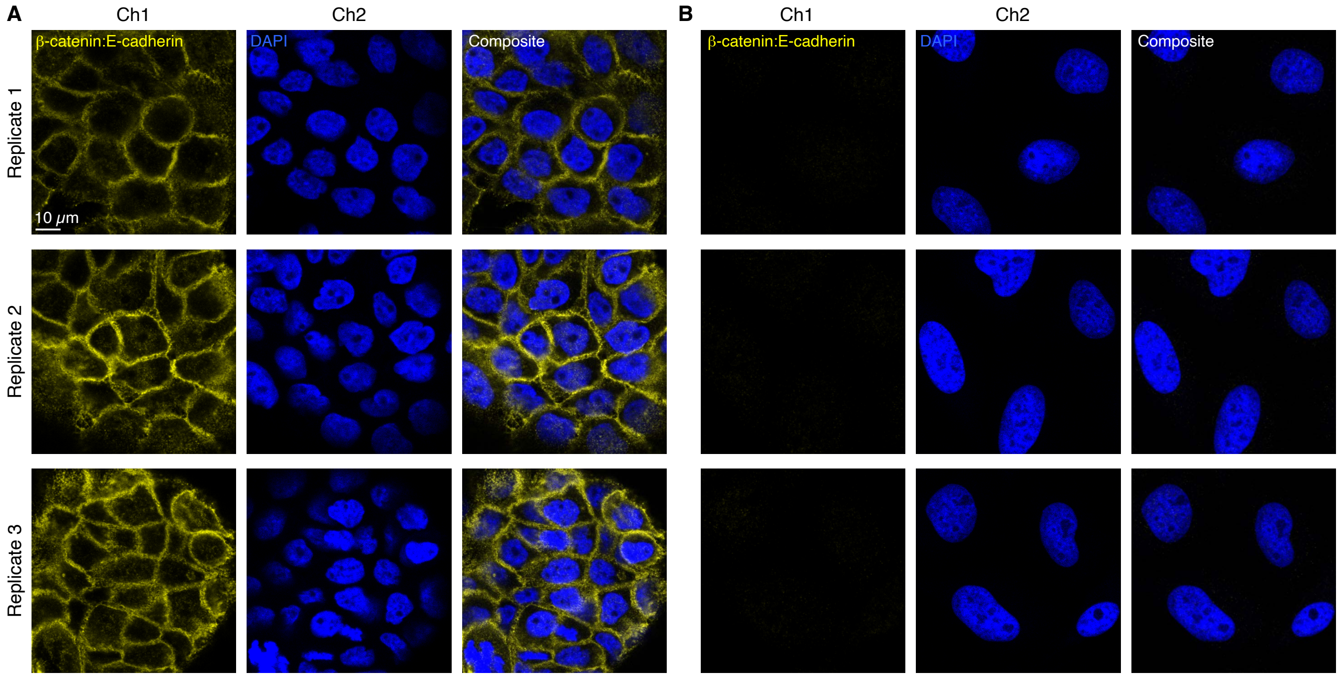

**Figure S2. Replicates for HCR imaging of a protein:protein complex in human cells (cf. Figure 3AB).** (A) A-431 cells. (B) HeLa cells. (A,B) 2-channel confocal images for 3 replicate wells on a multi-well coverslip; single optical section. Ch1: $\beta$-catenin:E-cadherin target complex (Alexa647). Ch2: DAPI.

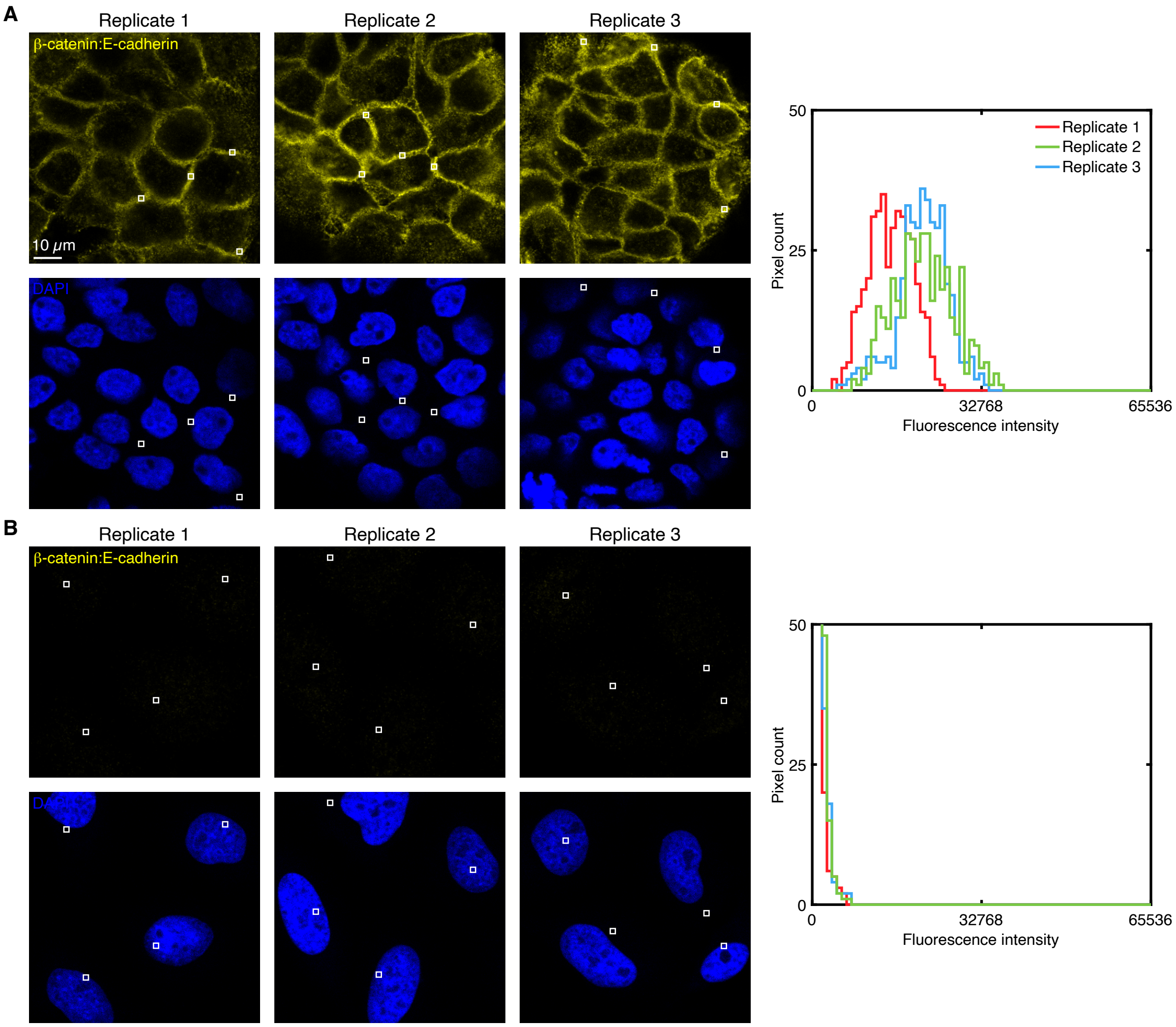

**Figure S3. Measurement of signal and background for protein:protein complex imaging in human cells (cf. Figure 3AB).** (A) Use experiment of Type 1a in Table S6 to measure SIG+BACK in a region of high protein:protein complex expression (pixels within rectangles) in 3 replicate wells of A-431 cells. (B) Use experiment of Type 1b in Table S6 to measure BACK in a region of maximum background (pixels within rectangles) in 3 replicate wells of HeLa cells. (A,B) Left: confocal images collected with the microscope settings optimized to avoid saturating SIG+BACK pixels; DAPI channel contextualizes placement of the rectangles; single optical section. Ch1: $\beta$-catenin:E-cadherin target complex (Alexa647). Ch2: DAPI. Right: pixel intensity histograms for rectangular regions of Ch1 (one rectangle in each of 4 individual cells in each of 3 replicate wells on a multi-well slide).

| Protein:protein complex | SIG+BACK | SIG | BACK | SIG/BACK |
|---|---|---|---|---|
| $\beta$-catenin:E-cadherin | 19 000 $\pm$ 2000 | 18 000 $\pm$ 2000 | 700 $\pm$ 70 | 26 $\pm$ 4 |

**Table S10. Estimated signal-to-background for protein:protein complex imaging in human cells (cf. Figure 3AB).** Estimated signal-to-background (SIG/BACK) based on the methods of Section S1.7.2 with SIG+BACK estimated in A-431 cells and BACK estimated in HeLa cells. Mean $\pm$ estimated standard error of the mean via uncertainty propagation for representative regions of $N = 3$ replicate wells on a multi-well slide. Analysis based on rectangular regions depicted in Figure S3.

#### S2.3 Replicates, signal, and background for protein:protein complex imaging in mouse proT cells (cf. Figure 3CD)

For the RUNX1:PU.1 target complex, Scid.adh.2C2 cells retrovirally transduced with a PU.1-expressing vector serve as the positive sample and provide an estimate of signal + background (SIG+BACK), while Scid.adh.2C2 cells retrovirally transduced with an empty vector serve as the negative sample and provide an estimate of background (BACK). Additional studies are presented as follows:

- Figure S4 displays images for $N = 3$ replicate wells on a multi-well slide for mouse proT cells retrovirally transduced with a PU.1 vector (panel A) or with an empty vector (panel B) (cf. Figure 3CD).
- Figure S5 displays representative regions used for measurement of signal and background.
- Table S11 displays estimated values for signal, background, and signal-to-background.

**Protocol:** Section S1.9.
**Sample:** Scid.adh.2C2 mouse proT cells retrovirally transduced with a PU.1-expressing vector or an empty vector.
**Reagents:** Table S1.

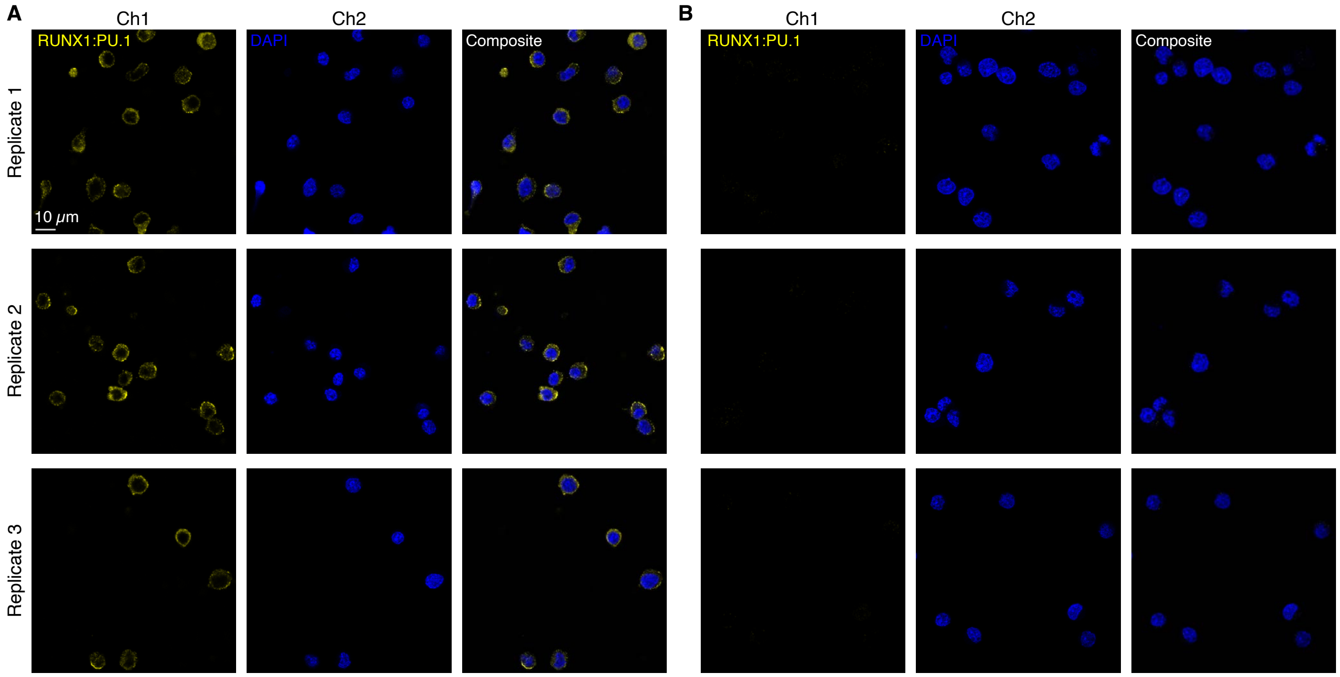

**Figure S4. Replicates for HCR imaging of a protein:protein complex in mouse proT cells (cf. Figure 3CD).** (A) Scid.adh.2C2 cells retrovirally transduced with a PU.1-expressing vector. (B) Scid.adh.2C2 cells retrovirally transduced with an empty vector. (A,B) 2-channel confocal images for 3 replicate wells on a multi-well slide; single optical section. Ch1: RUNX1:PU.1 target complex (Alexa647). Ch2: DAPI.

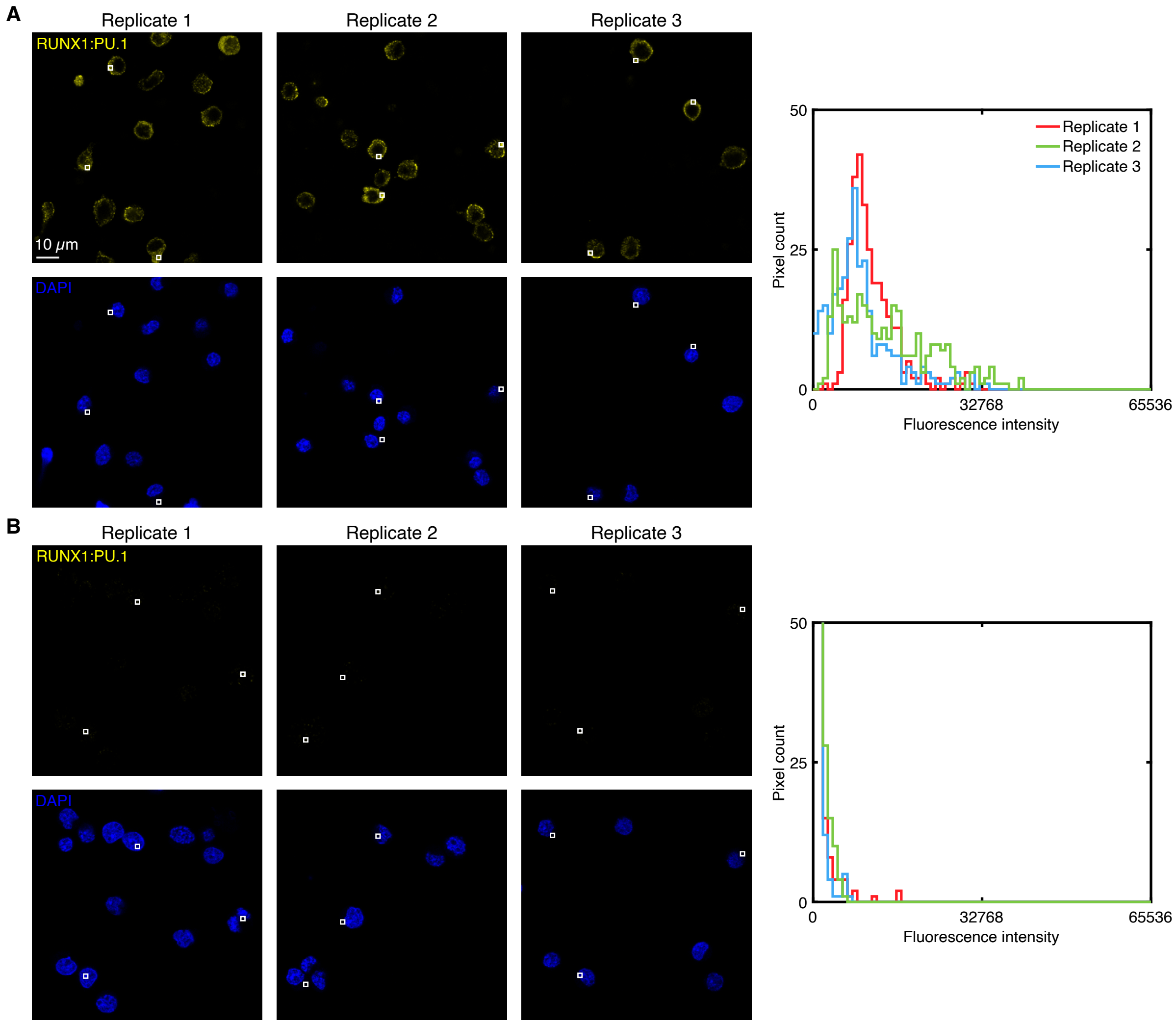

**Figure S5. Measurement of signal and background for protein:protein complex imaging in mouse proT cells (cf. Figure 3CD).** (A) Use experiment of Type 1a in Table S6 to measure SIG+BACK in a region of high protein:protein complex expression (pixels within rectangles) in 3 replicate wells of Scid.adh.2C2 retrovirally transduced with a PU.1-expressing vector. (B) Use experiment of Type 1b in Table S6 to measure BACK in a region of maximum background (pixels within rectangles) in 3 replicate wells of fixed Scid.adh.2C2 retrovirally transduced with an empty vector. (A,B) Left: confocal images collected with the microscope settings optimized to avoid saturating SIG+BACK pixels; DAPI channel contextualizes placement of the rectangles; single optical section. Ch1: RUNX1:PU.1 target complex (Alexa647). Ch2: DAPI. Right: pixel intensity histograms for rectangular regions of Ch1 (one rectangle in each of 3 individual cells in each of 3 replicate wells on a multi-well slide).

| Protein:protein complex | SIG+BACK | SIG | BACK | SIG/BACK |
|---|---|---|---|---|
| RUNX1:PU.1 | $11\,000 \pm 1000$ | $11\,000 \pm 1000$ | $700 \pm 100$ | $15 \pm 3$ |

**Table S11. Estimated signal-to-background for protein:protein complex imaging in mouse proT cells (cf. Figure 3CD).** Estimated signal-to-background (SIG/BACK) based on the methods of Section S1.7.2 with SIG+BACK estimated in Scid.adh.2C2 cells retrovirally transduced with a PU.1-expressing vector and BACK estimated in Scid.adh.2C2 cells retrovirally transduced with an empty vector. Mean $\pm$ estimated standard error of the mean via uncertainty propagation for representative regions of $N = 3$ replicate wells on a multi-well slide. Analysis based on rectangular regions depicted in Figure S5.

#### S2.4 Replicates, signal, and background for protein:protein complex imaging in FFPE human breast tissue sections (cf. Figure 3EF)

For the $\beta$-catenin:E-cadherin target complex, FFPE normal human breast tissue sections serve as the positive sample and provide an estimate of signal + background (SIG+BACK), while FFPE invasive lobular carcinoma human breast tissue sections from the same patient serve as the negative sample and provide an estimate of background (BACK). Additional studies are presented as follows:

- Figure S6 displays images for $N = 3$ replicate FFPE human breast tissue sections for normal (panel A) and invasive lobular carcinoma (panel B) (cf. Figure 3EF).
- Figure S7 displays representative regions used for measurement of signal and background.
- Table S12 displays estimated values for signal, background, and signal-to-background.

**Protocol:** Section S1.10.
**Sample:** FFPE human breast tissue sections (normal or invasive lobular carcinoma); thickness: 5 $\mu$m.
**Reagents:** Table S1.

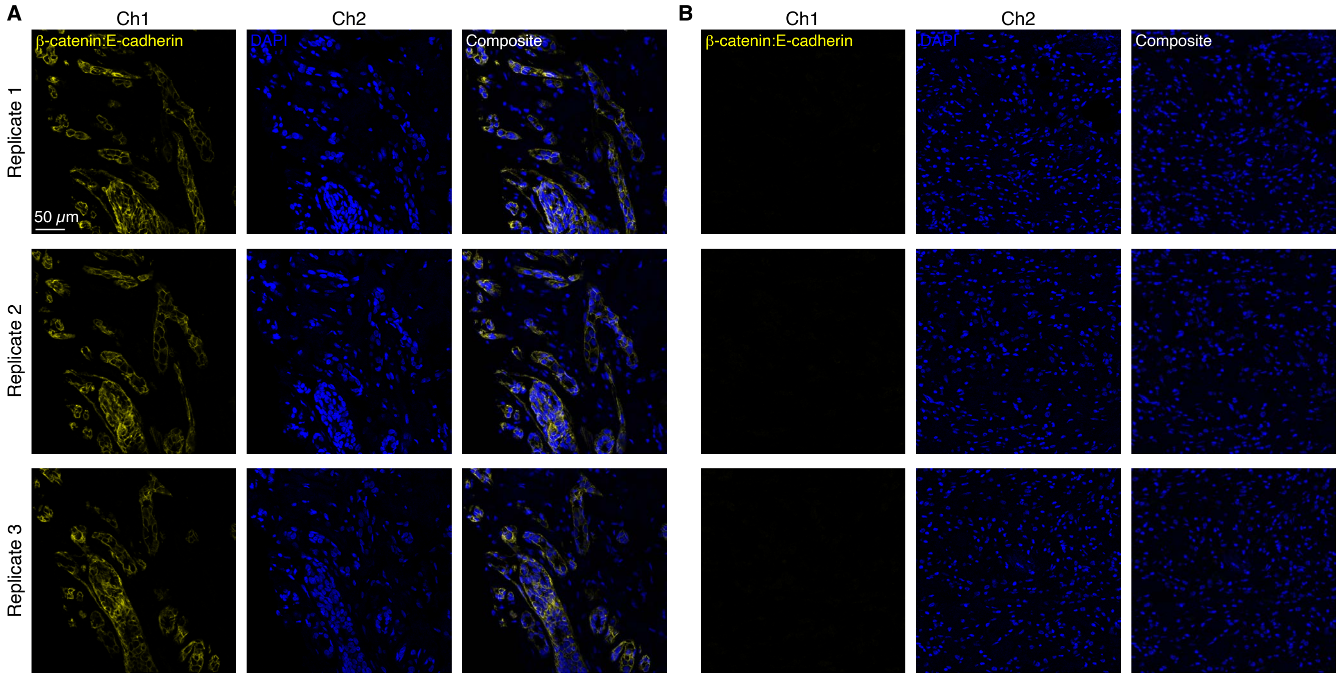

**Figure S6. Replicates for HCR imaging of a protein:protein complex in FFPE human breast tissue sections (cf. Figure 3EF).** (A) FFPE normal human breast tissue sections. (B) FFPE invasive lobular carcinoma human breast tissue sections. (A,B) 2-channel confocal images for 3 replicate FFPE human breast tissue sections; single optical section. Ch1: $\beta$-catenin:E-cadherin target complex (Alexa647). Ch2: DAPI.

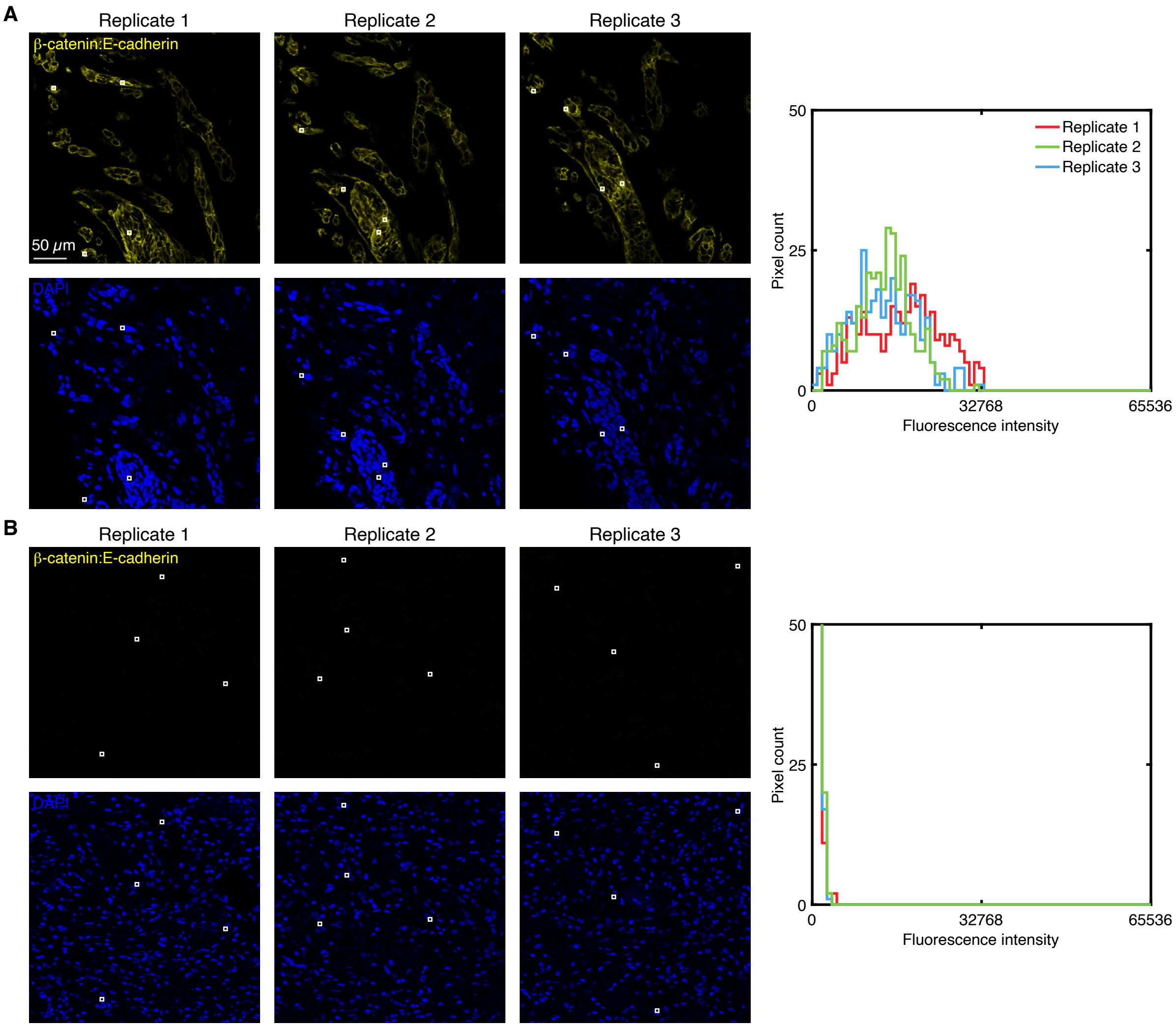

**Figure S7. Measurement of signal and background for protein:protein complex imaging in FFPE human breast tissue sections (cf. Figure 3EF).** (A) Use experiment of Type 1a in Table S6 to measure SIG+BACK in a region of high protein:protein complex expression (pixels within rectangles) in 3 replicate FFPE normal human breast tissue sections. (B) Use experiment of Type 1b in Table S6 to measure BACK in a region of maximum background (pixels within rectangles) in 3 replicate FFPE invasive lobular carcinoma human breast tissue sections. (A,B) Left: confocal images collected with the microscope settings optimized to avoid saturating SIG+BACK pixels; DAPI channel contextualizes placement of the rectangles; single optical section. Ch1: $\beta$-catenin:E-cadherin target complex (Alexa647). Ch2: DAPI. Right: pixel intensity histograms for rectangular regions of Ch1 (4 rectangles in each of 3 replicate FFPE human breast tissue sections).

| Protein:protein complex | SIG+BACK | SIG | BACK | SIG/BACK |
|---|---|---|---|---|
| $\beta$-catenin:E-cadherin | $14\,000 \pm 1000$ | $14\,000 \pm 1000$ | $460 \pm 8$ | $30 \pm 3$ |

**Table S12. Estimated signal-to-background for protein:protein complex imaging in FFPE human breast tissue sections (cf. Figure 3EF).** Estimated signal-to-background (SIG/BACK) based on the methods of Section S1.7.2 with SIG+BACK estimated in FFPE normal human breast tissue sections and BACK estimated in FFPE invasive lobular carcinoma human breast tissue sections. Mean $\pm$ estimated standard error of the mean via uncertainty propagation for representative regions in $N = 3$ replicate tissue sections. Analysis based on rectangular regions depicted in Figure S7.

#### S2.5 Replicates, signal, and background for 3-plex protein:protein complex imaging in human cells (cf. Figure 4)

For 3-plex protein:protein complex imaging using HCR in human cells on a multi-well slide, the 4 channels are (3 protein:protein complexes + DAPI):

- **Ch1:** Protein:protein complex $\alpha$-tubulin:$\beta$-tubulin, amplifier B6-Alexa488.
- **Ch2:** Protein:protein complex $\beta$-catenin:E-cadherin, amplifier B1-Alexa546.
- **Ch3:** Protein:protein complex SC35:SON, amplifier B9-Alexa647.
- **Ch4:** DAPI.

Additional studies are presented as follows:

- Figure S8 displays 3-plex images for $N = 3$ replicate wells on a multi-well slide (cf. Figure 4).
- Figures S9–S11 display representative regions of individual channels used for measurement of signal and background for each target.
- Table S13 displays estimated values for signal, background, and signal-to-background for each channel.

**Protocol:** Section S1.8.
**Sample:** A-431 cells.
**Reagents:** Table S1.

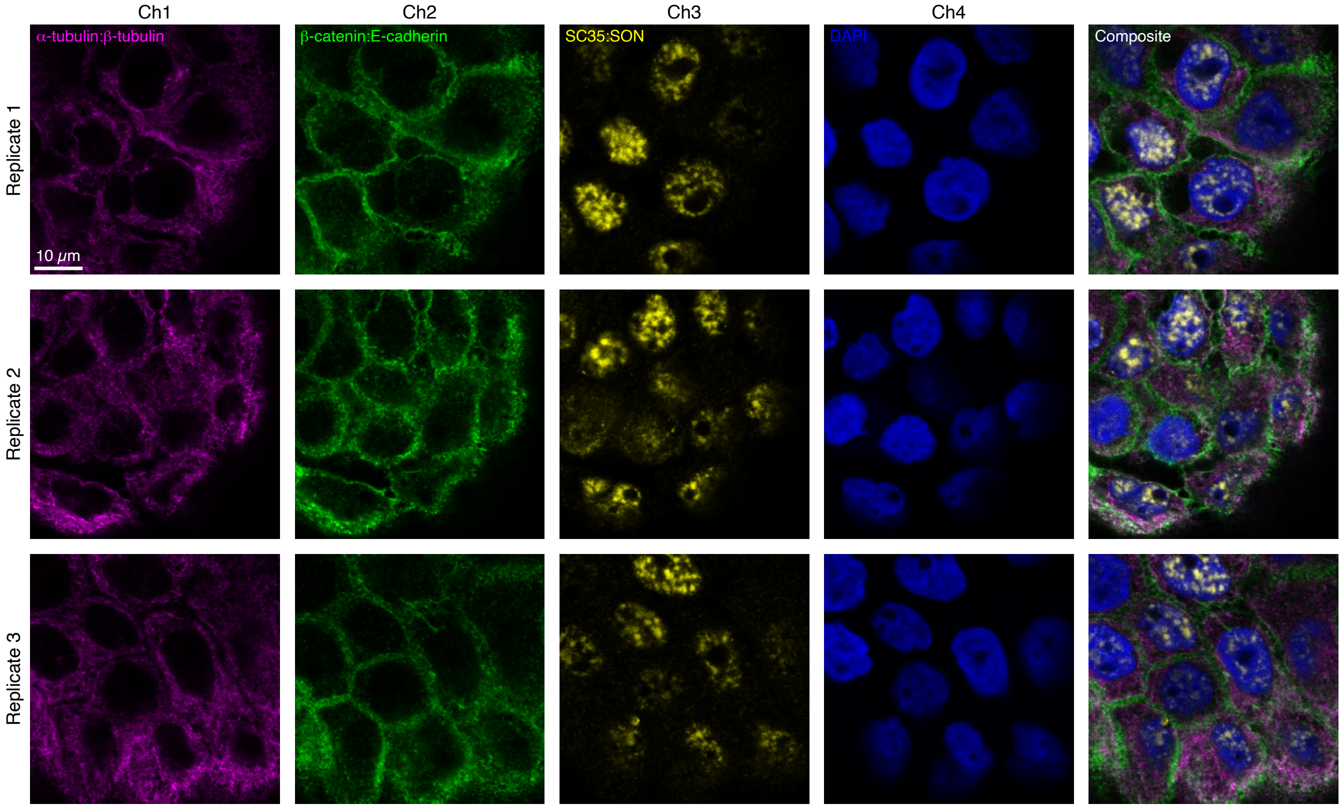

**Figure S8. Replicates for 3-plex protein:protein complex imaging using HCR in human cells (cf. Figure 4).** 4-channel images for 3 replicate wells on a multi-well slide; single optical section. Ch1: $\alpha$-tubulin:$\beta$-tubulin target complex (Alexa488). Ch2: $\beta$-catenin:E-cadherin target complex (Alexa546). Ch3: SC35:SON target complex (Alexa647). Ch4: DAPI. Sample: A-431 cells.

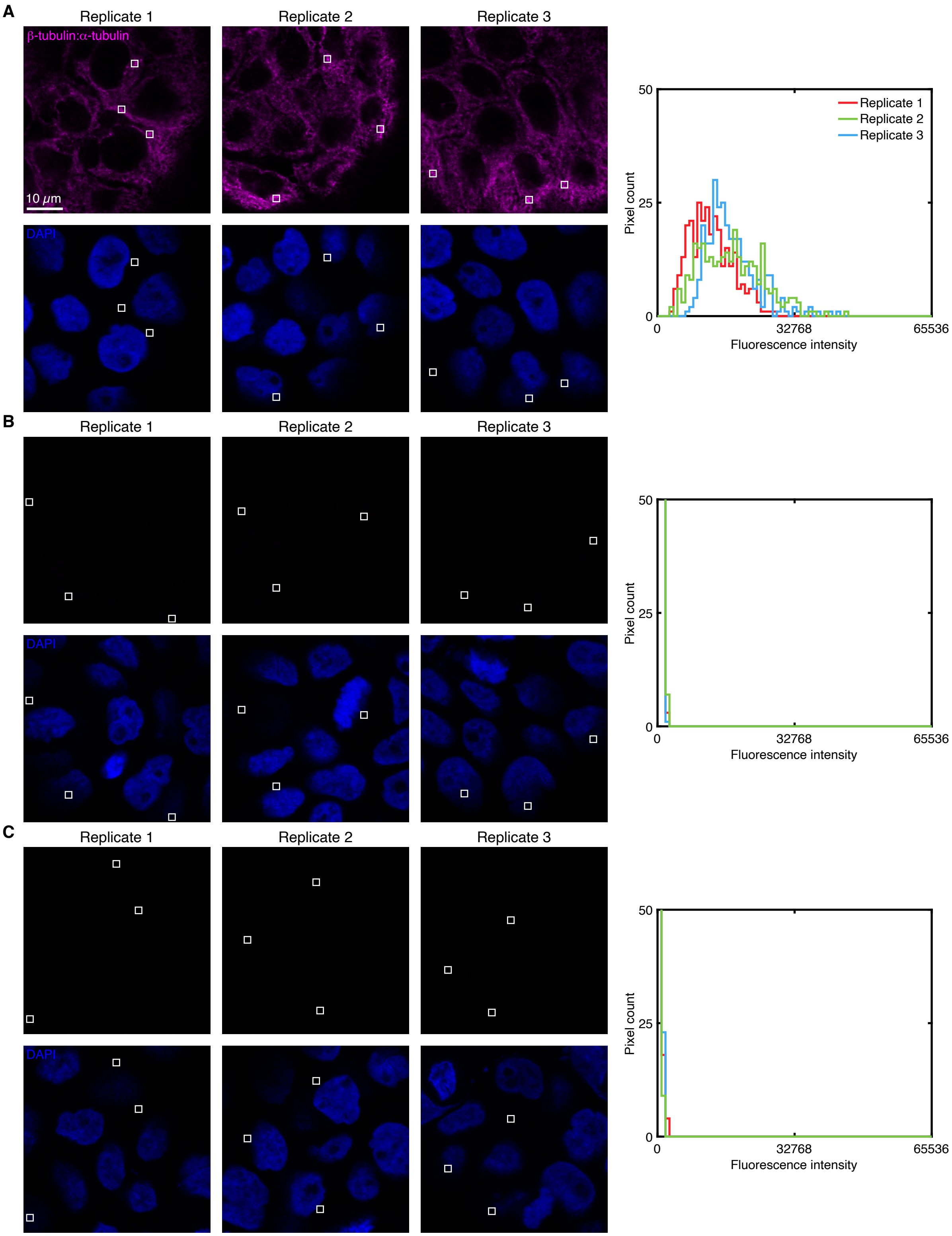

**Figure S9. Measurement of signal and background for protein:protein complex $\alpha$-tubulin:$\beta$-tubulin in human cells (cf. Figure 4).** (A) Use experiment of Type 1a in Table S6 to measure SIG+BACK in a region of high protein:protein complex expression (pixels within rectangles) in 3 replicate wells. (B) Use experiment of Type 2a in Table S6 to measure $NSD_{P1}$+NSA+AF in a region of maximum background (pixels within rectangles) in 3 replicate wells. (C) Use experiment of Type 2b in Table S6 to measure $NSD_{P2}$+NSA+AF in a region of maximum background (pixels within rectangles) in 3 replicate wells. (A,B,C) Left: confocal images collected with the microscope settings optimized to avoid saturating SIG+BACK pixels; DAPI channel contextualizes placement of the rectangles; single optical section. Ch1: $\alpha$-tubulin:$\beta$-tubulin target complex (Alexa488). Ch4: DAPI. Right: pixel intensity histograms for rectangular regions of Ch1 (one rectangle in each of 3 individual cells in each of 3 replicate wells on a multi-well slide).

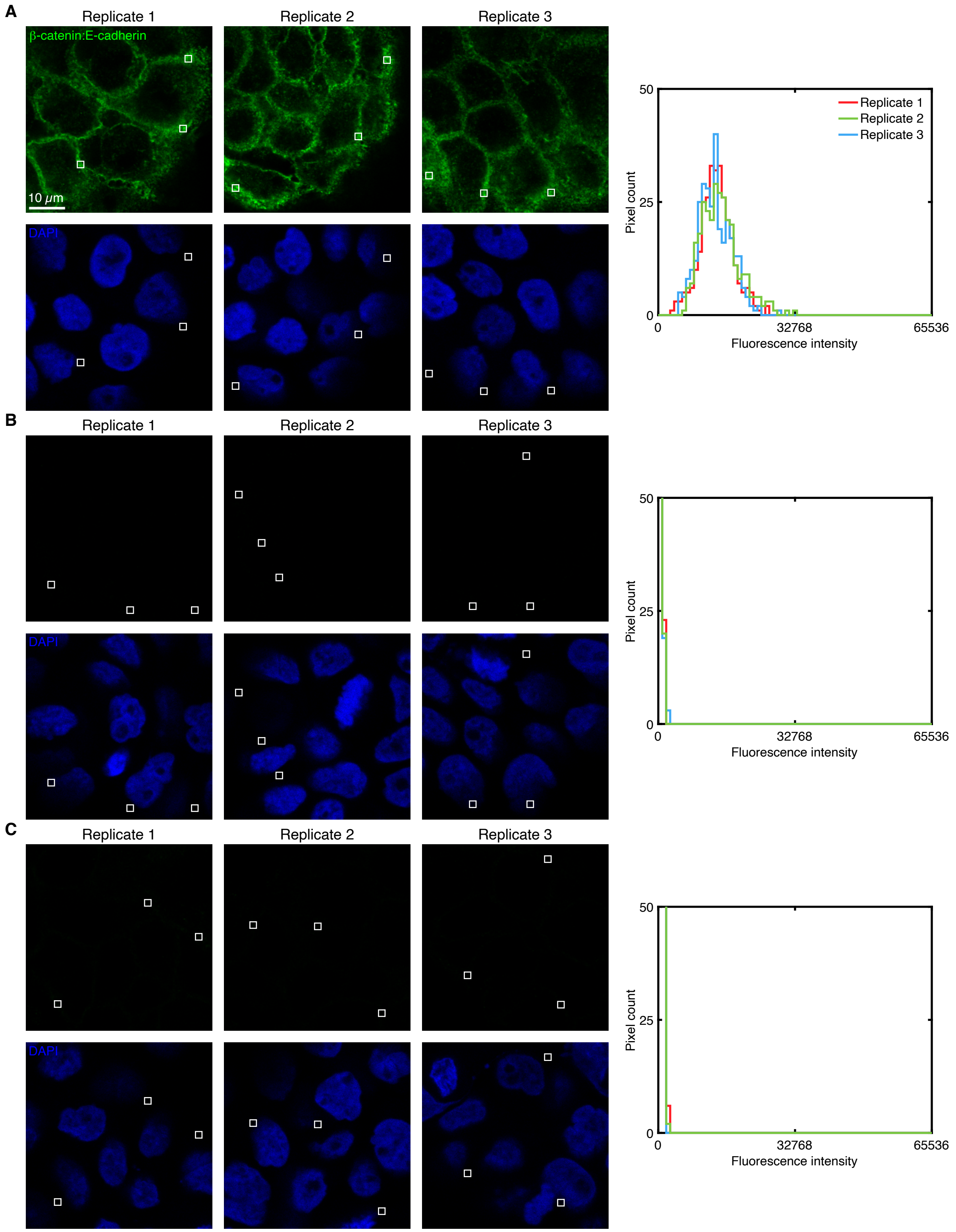

**Figure S10. Measurement of signal and background for protein:protein complex $\beta$-catenin:E-cadherin in human cells (cf. Figure 4).** (A) Use experiment of Type 1a in Table S6 to measure SIG+BACK in a region of high protein:protein complex expression (pixels within rectangles) in 3 replicate wells. (B) Use experiment of Type 2a in Table S6 to measure $NSD_{P1}$+NSA+AF in a region of maximum background (pixels within rectangles) in 3 replicate wells. (C) Use experiment of Type 2b in Table S6 to measure $NSD_{P2}$+NSA+AF in a region of maximum background (pixels within rectangles) in 3 replicate wells. (A,B,C) Left: confocal images collected with the microscope settings optimized to avoid saturating SIG+BACK pixels; DAPI channel contextualizes placement of the rectangles; single optical section. Ch2: $\beta$-catenin:E-cadherin target complex (Alexa546). Ch4: DAPI. Right: pixel intensity histograms for rectangular regions of Ch2 (one rectangle in each of 3 individual cells in each of 3 replicate wells on a multi-well coverslip).

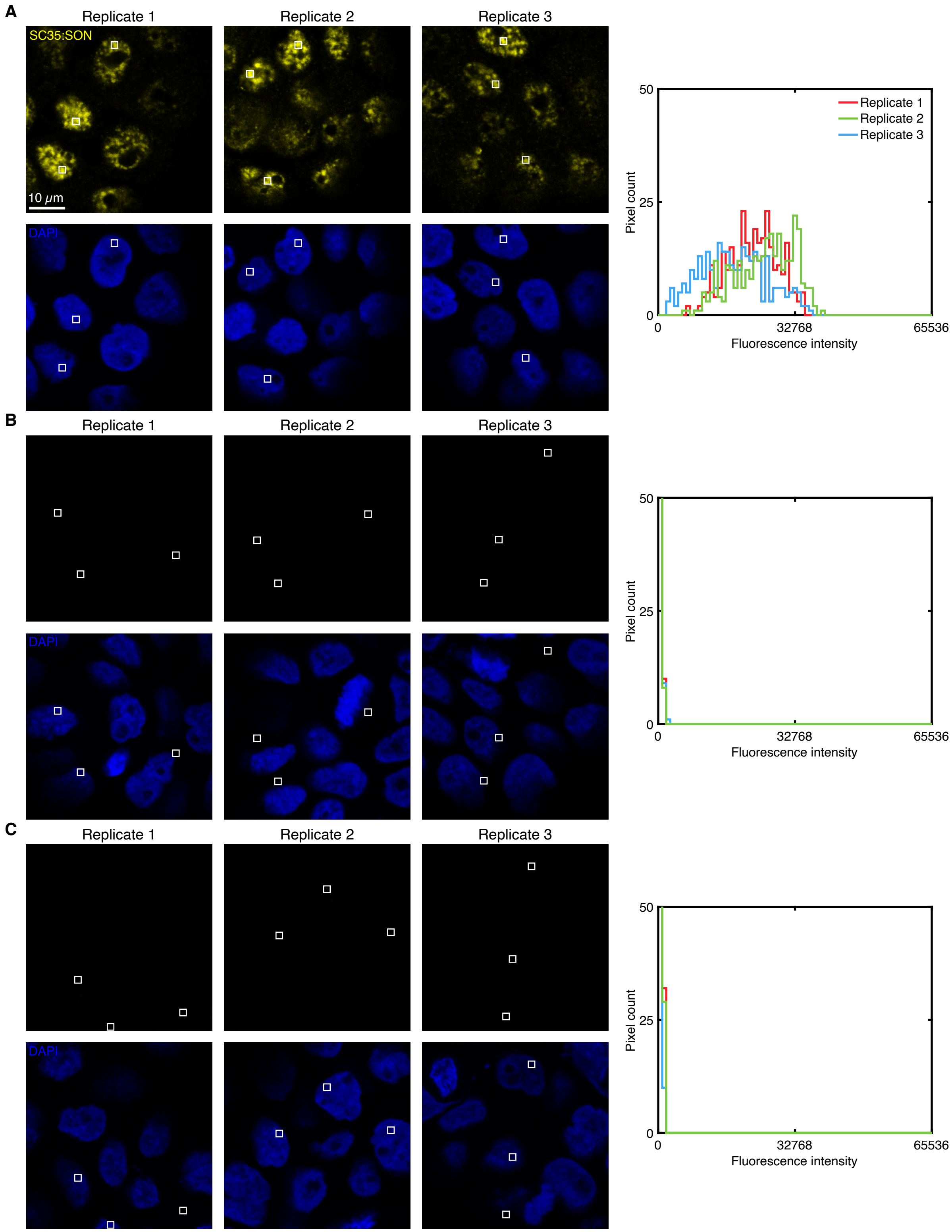

**Figure S11. Measurement of signal and background for protein:protein complex SC35:SON in human cells (cf. Figure 4).** (A) Use experiment of Type 1a in Table S6 to measure SIG+BACK in a region of high protein:protein complex expression (pixels within rectangles) in 3 replicate wells. (B) Use experiment of Type 2a in Table S6 to measure $NSD_{P1}$+NSA+AF in a region of maximum background (pixels within rectangles) in 3 replicate wells. (C) Use experiment of Type 2b in Table S6 to measure $NSD_{P2}$+NSA+AF in a region of maximum background (pixels within rectangles) in 3 replicate wells. (A,B,C) Left: confocal images collected with the microscope settings optimized to avoid saturating SIG+BACK pixels; DAPI channel contextualizes placement of the rectangles; single optical section. Ch3: SC35:SON target complex (Alexa647). Ch4: DAPI. Right: pixel intensity histograms for rectangular regions of Ch3 (one rectangle in each of 3 individual cells in each of 3 replicate wells on a multi-well coverslip).

| Protein:protein complex | SIG+BACK | $NSD_{P1}$+NSA+AF | $NSD_{P2}$+NSA+AF | BACK | SIG | SIG/BACK |
|---|---|---|---|---|---|---|
| $\alpha$-tubulin:$\beta$-tubulin | $16\,000 \pm 2000$ | $350 \pm 20$ | $130 \pm 20$ | $480 \pm 30$ | $15\,000 \pm 2000$ | $32 \pm 4$ |
| $\beta$-catenin:E-cadherin | $13\,700 \pm 400$ | $168 \pm 8$ | $430 \pm 90$ | $600 \pm 90$ | $13\,100 \pm 400$ | $22 \pm 4$ |
| SC35:SON | $22\,000 \pm 2000$ | $86 \pm 4$ | $140 \pm 30$ | $230 \pm 30$ | $21\,000 \pm 2000$ | $90 \pm 20$ |

**Table S13. Estimated signal-to-background for 3-plex protein:protein complex imaging in human cells (cf. Figure 4).** Estimated signal-to-background (SIG/BACK) based on the methods of Section S1.7.2 with SIG+BACK estimated using an experiment of Type 1a in Table S6. BACK is estimated as the sum of $NSD_{P1}$+NSA+AF and $NSD_{P2}$+NSA+AF obtained from experiments of Type 2a and 2b in Table S6. Mean $\pm$ estimated standard error of the mean via uncertainty propagation for representative regions of $N = 3$ replicate wells on a multi-well slide. Analysis based on rectangular regions depicted in Figures S9–S11.

#### S2.6 qHCR imaging: protein:protein complex relative quantitation with subcellular resolution in an anatomical context (cf. Figure 5)

##### S2.6.1 Redundant 2-channel imaging of the $\beta$-catenin:E-cadherin target complex in human cells

Here, we perform redundant 2-channel imaging of the $\beta$-catenin:E-cadherin target complex in human cells. Each protein is detected by a 1°Ab probe, which is then redundantly detected by two batches of split-initiator 2°Ab probes that interact with orthogonal proximity probes to colocalize full HCR initiators that trigger orthogonal spectrally distinct HCR amplifiers.

Additional studies are presented as follows:

- Figure S12 displays 2-plex images and 2-channel voxel intensity scatter plots for $\beta$-catenin:E-cadherin target complex in fixed A-431 cells for $N = 3$ replicate wells on a multi-well slide.
- Figures S13 and S14 display representative regions used for determining the BOT values used to normalize data for Figure S12C using methods of Section S1.7.3.
- Table S14 displays values used for signal normalization in Figure S12.
- Figures S15 and S16 display representative regions used for measurement of signal and background for the $\beta$-catenin:E-cadherin target complex.
- Table S15 displays estimated values for signal, background, and signal-to-background for each channel.

**Protocol:** Section S1.10.
**Sample:** A-431 cells.
**Reagents:** Table S1.

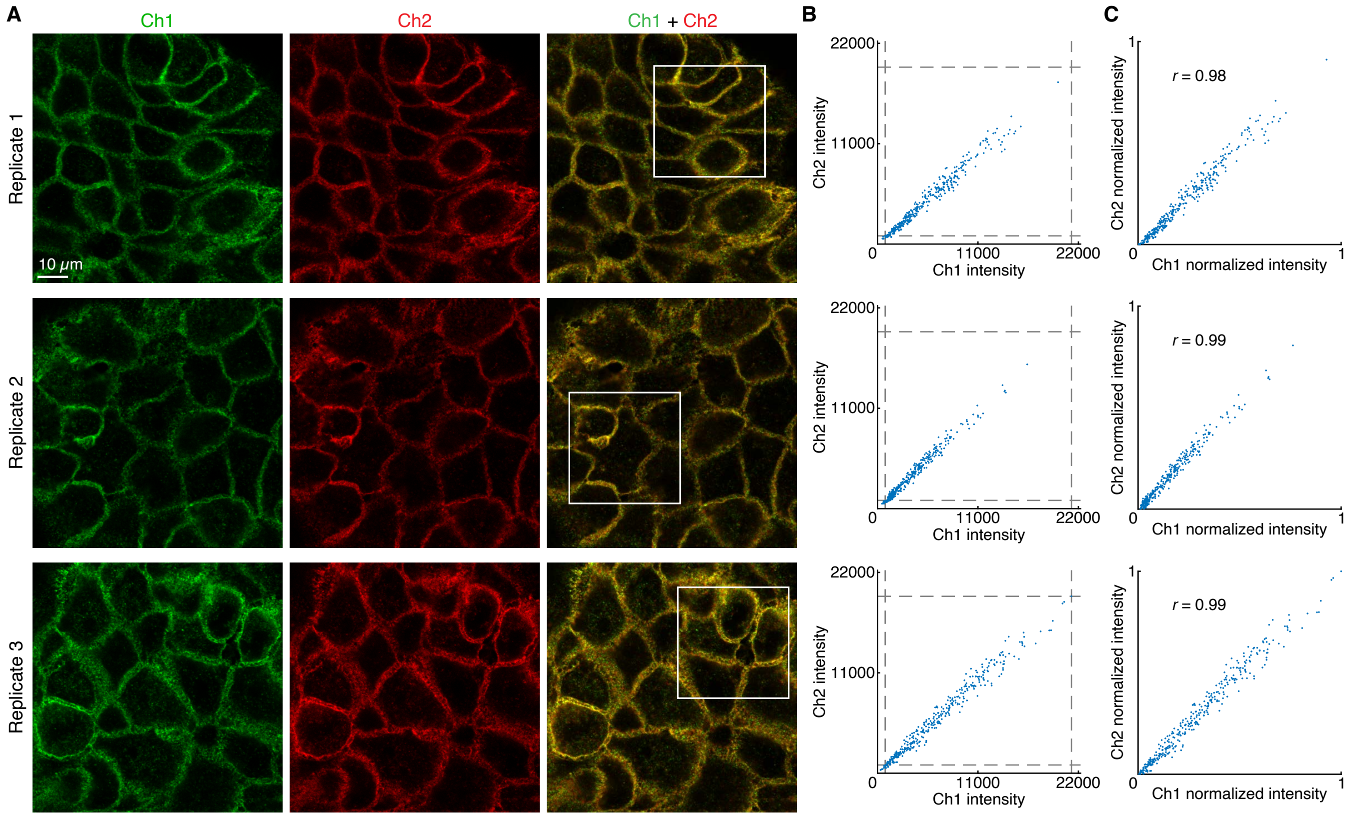

**Figure S12. Redundant 2-channel detection of $\beta$-catenin:E-cadherin target complex in human cells (cf. Figure 5).** (A) Confocal images: individual channels and merge; single optical section. Ch1: $\beta$-catenin:E-cadherin (Alexa546). Ch2: $\beta$-catenin:E-cadherin (Alexa647). Solid boundaries denote regions of variable expression. Pixel size: $0.18\times0.18\times0.8$ $\mu$m. Sample: Fixed A-431 cells. (B) Raw voxel intensity scatter plots representing signal plus background for voxels within solid boundaries of panel A. Voxel size: $2.0\times2.0\times0.8$ $\mu$m. Dashed lines represent BOT and TOP values (Table S14) used to normalize data for panel C using methods of Section S1.7.3. The BOT value for each channel was determined from the solid boundaries (regions of no/low protein:protein complex expression) in Figures S13 (Ch 1) and S14 (Ch 2). (C) Normalized voxel intensity scatter plots representing normalized signal (Pearson correlation coefficient, $r$).

| Channel | Protein:protein complex | Fluorophore | BOT | TOP |
|---|---|---|---|---|
| Ch1 | $\beta$-catenin:E-cadherin | Alexa546 | 837 | 21241 |
| Ch2 | $\beta$-catenin:E-cadherin | Alexa647 | 901 | 19384 |

**Table S14. BOT and TOP values used to calculate normalized voxel intensities for scatter plots of Figures 6C (top panel) and S12C using methods of Section S1.7.3.** Analysis based on rectangular regions depicted in Figures S12A (Ch1: TOP; Ch2: TOP), S13 (Ch 1: BOT) and S14 (Ch2: BOT). BOT calculated based on background estimate BACK $\approx$ $\mathrm{NSD_{P1}}$+NSA+AF + $\mathrm{NSD_{P2}}$+NSA+AF using methods of Section S1.7.3.

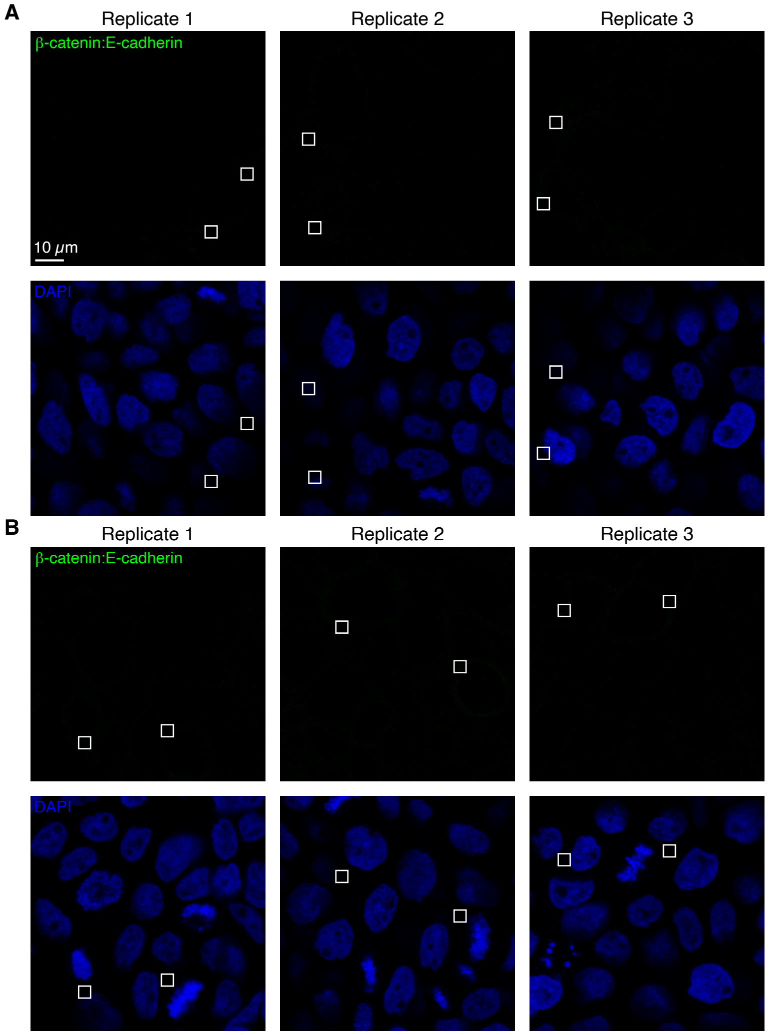

**Figure S13. Measurement of BOT for Ch1 for redundant 2-channel detection of $\beta$-catenin:E-cadherin target complex in human cells (cf. Figure 5).** (A) Experiment of Type 2a in Table S6 to measure $NSD_{P1}$+NSA+AF in a region of maximum background. (B) Experiment of Type 2b in Table S6 to measure $NSD_{P2}$+NSA+AF in a region of maximum background. (A,B) Solid boundaries denote representative regions of maximum background. BOT calculated based on background estimate BACK $\approx$ $NSD_{P1}$+NSA+AF + $NSD_{P2}$+NSA+AF using methods of Section S1.7.3. Confocal images; single optical section. Sample: A-431 cells.

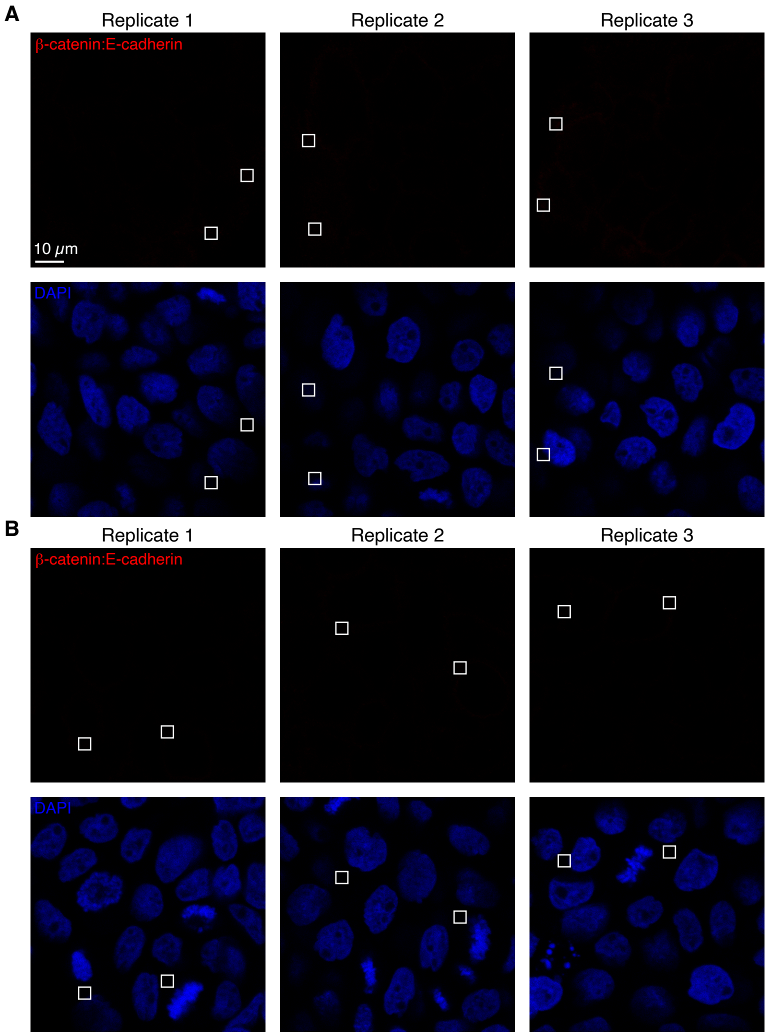

**Figure S14. Measurement of BOT for Ch2 for redundant 2-channel detection of $\beta$-catenin:E-cadherin target complex in human cells (cf. Figure 5).** (A) Experiment of Type 2a in Table S6 to measure $\text{NSD}_{\text{P1}}$+NSA+AF in a region of maximum background. (B) Experiment of Type 2b in Table S6 to measure $\text{NSD}_{\text{P2}}$+NSA+AF in a region of maximum background. (A,B) Solid boundaries denote representative regions of maximum background. BOT calculated based on background estimate BACK $\approx$ $\text{NSD}_{\text{P1}}$+NSA+AF + $\text{NSD}_{\text{P2}}$+NSA+AF using methods of Section S1.7.3. Confocal images; single optical section. Sample: A-431 cells.

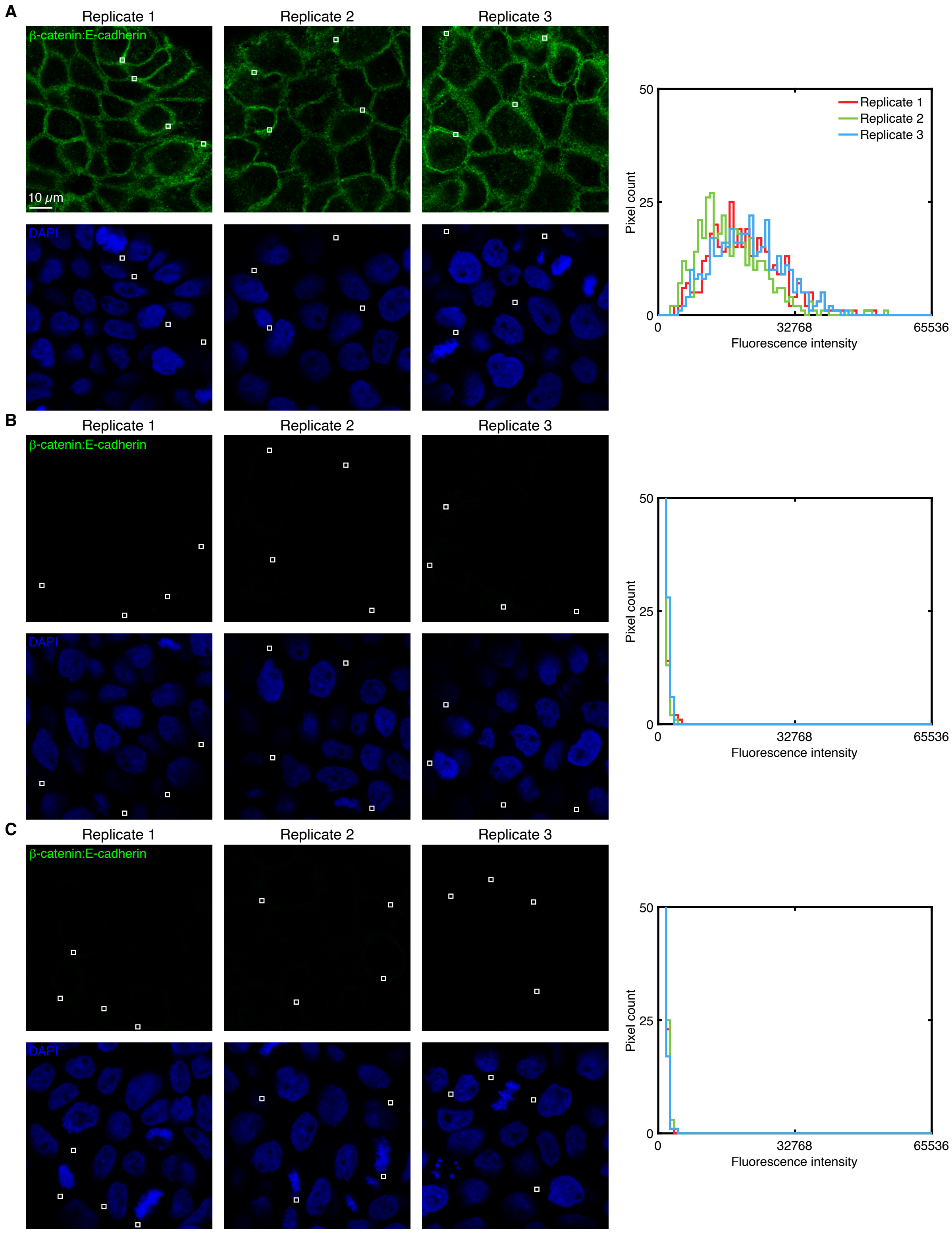

**Figure S15. Measurement of Ch1 signal and background for redundant 2-channel detection of protein:protein complex $\beta$-catenin:E-cadherin in fixed A-431 cells (cf. Figure 5).** (A) Ch1 (Alexa546) for redundant 2-channel imaging of $\beta$-catenin:E-cadherin target complex using an experiment of Type 1a in Table S6 to measure SIG + BACK. Solid boundaries denote regions of high expression. (B) Experiment of Type 2a in Table S6 to measure $NSD_{P1}$+NSA+AF in a region of maximum background. (C) Experiment of Type 2b in Table S6 to measure $NSD_{P2}$+NSA+AF in a region of maximum background. (B,C) Solid boundaries denote representative regions of maximum background. (A,B,C) Left: confocal images; single optical section. Right: pixel intensity histograms for the depicted regions. Sample: A-431 cells.

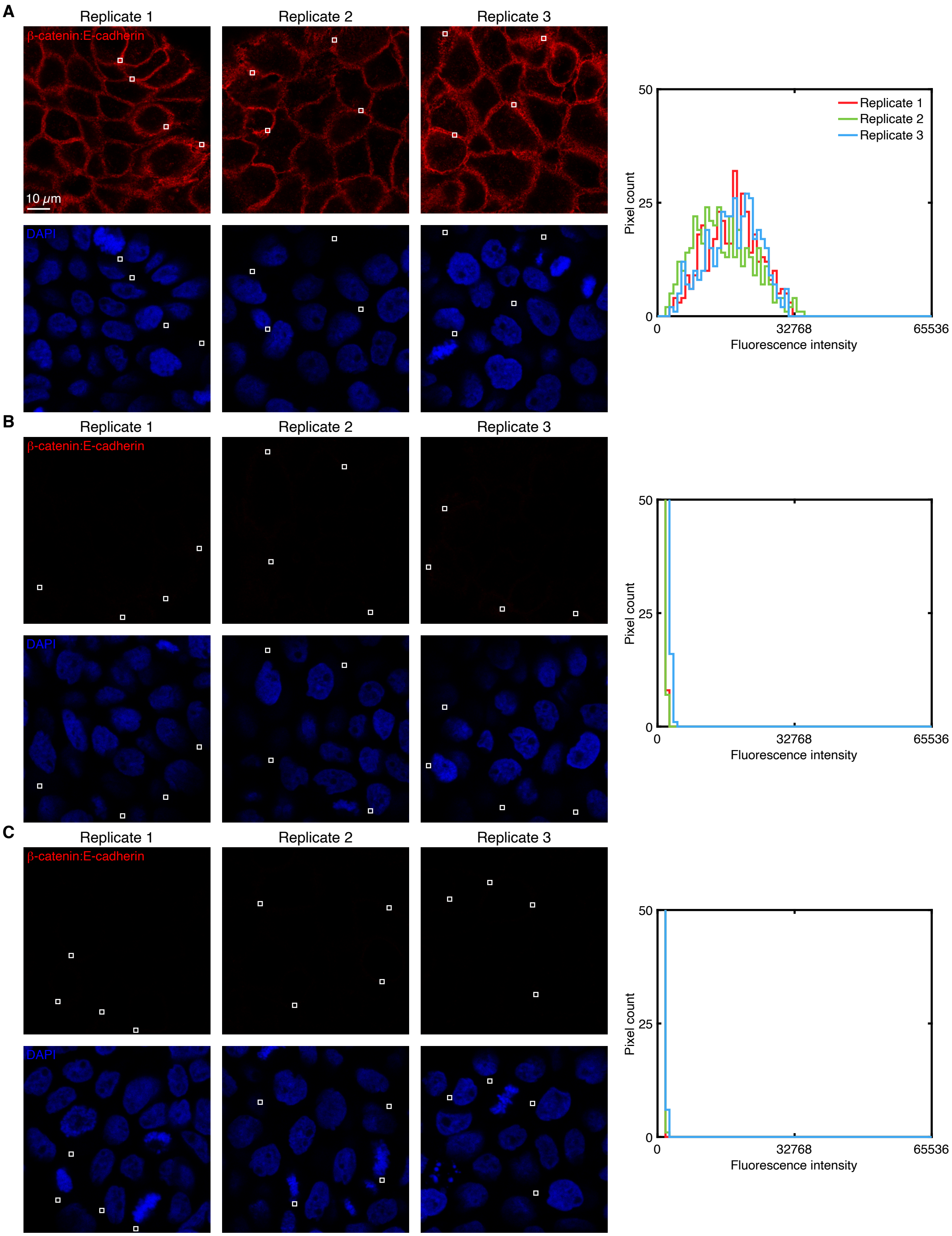

**Figure S16. Measurement of Ch2 signal and background for redundant 2-channel detection of $\beta$-catenin:E-cadherin target complex in human cells (cf. Figure 5).** (A) Ch2 (Alexa647) for redundant 2-channel imaging of $\beta$-catenin:E-cadherin target complex using an experiment of Type 1a in Table S6 to measure SIG + BACK. Solid boundaries denote regions of high expression. (B) Experiment of Type 2a in Table S6 to measure $NSD_{P1}$+NSA+AF in a region of maximum background. Solid boundaries denote regions of representative regions of maximum background. (C) Experiment of Type 2b in Table S6 to measure $NSD_{P2}$+NSA+AF in a region of maximum background. (B,C) Solid boundaries denote representative regions of maximum background. (A,B,C) Left: confocal images; single optical section. Right: pixel intensity histograms for the depicted regions. Sample: A-431 cells.

| Channel | Protein:protein complex | Fluorophore | $\mathrm{NSD_{P1}}$+NSA+AF | $\mathrm{NSD_{P2}}$+NSA+AF | BACK | SIG+BACK | SIG | SIG/BACK |
|---|---|---|---|---|---|---|---|---|
| Ch1 | $\beta$-catenin:E-cadherin | Alexa546 | $490 \pm 70$ | $640 \pm 20$ | $1130 \pm 80$ | $20\,000 \pm 1000$ | $18\,000 \pm 1000$ | $16 \pm 2$ |
| Ch2 | $\beta$-catenin:E-cadherin | Alexa647 | $700 \pm 200$ | $380 \pm 10$ | $1000 \pm 200$ | $16\,700 \pm 800$ | $15\,700 \pm 800$ | $15 \pm 3$ |

**Table S15. Estimated signal-to-background for redundant 2-channel detection of $\beta$-catenin:E-cadherin target complex in human cells (cf. Figure 5).** SIG + BACK is measured using an experiment of Type 1a in Table S6. BACK is measured using experiments of Type 2a and Type 2b in Table S6 using the estimate BACK $\approx$ $\mathrm{NSD_{P1}}$+NSA+AF + $\mathrm{NSD_{P2}}$+NSA+AF. Mean $\pm$ estimated standard error of the mean via uncertainty propagation for $N = 3$ replicate wells on a multi-well slide. Analysis based on rectangular regions depicted in Figures S15 and S16 using methods of Section S1.7.2.

##### S2.6.2 Redundant 2-channel imaging of the $\beta$-catenin:E-cadherin target complex in FFPE human breast tissue sections

Here, we perform redundant 2-channel imaging of the $\beta$-catenin:E-cadherin target complex in FFPE human breast tissue sections. Each protein is detected by a 1°Ab probe, which is then redundantly detected by two batches of split-initiator 2°Ab probes that interact with orthogonal proximity probes to colocalize full HCR initiators that trigger orthogonal spectrally distinct HCR amplifiers.

Additional studies are presented as follows:

- Figure S17 displays 2-plex images and 2-channel voxel intensity scatter plots for $\beta$-catenin:E-cadherin target complex in $N = 3$ replicate FFPE human breast tissue sections.
- Table S16 displays values used for signal normalization in Figures S17.
- Figure S18 displays representative regions used for measurement of signal and background for $\beta$-catenin:E-cadherin target complex.
- Table S17 displays estimated values for signal, background, and signal-to-background for each channel.

**Protocol:** Section S1.10.
**Sample:** FFPE normal human breast tissue sections; thickness: 5 $\mu$m.
**Reagents:** Table S1.

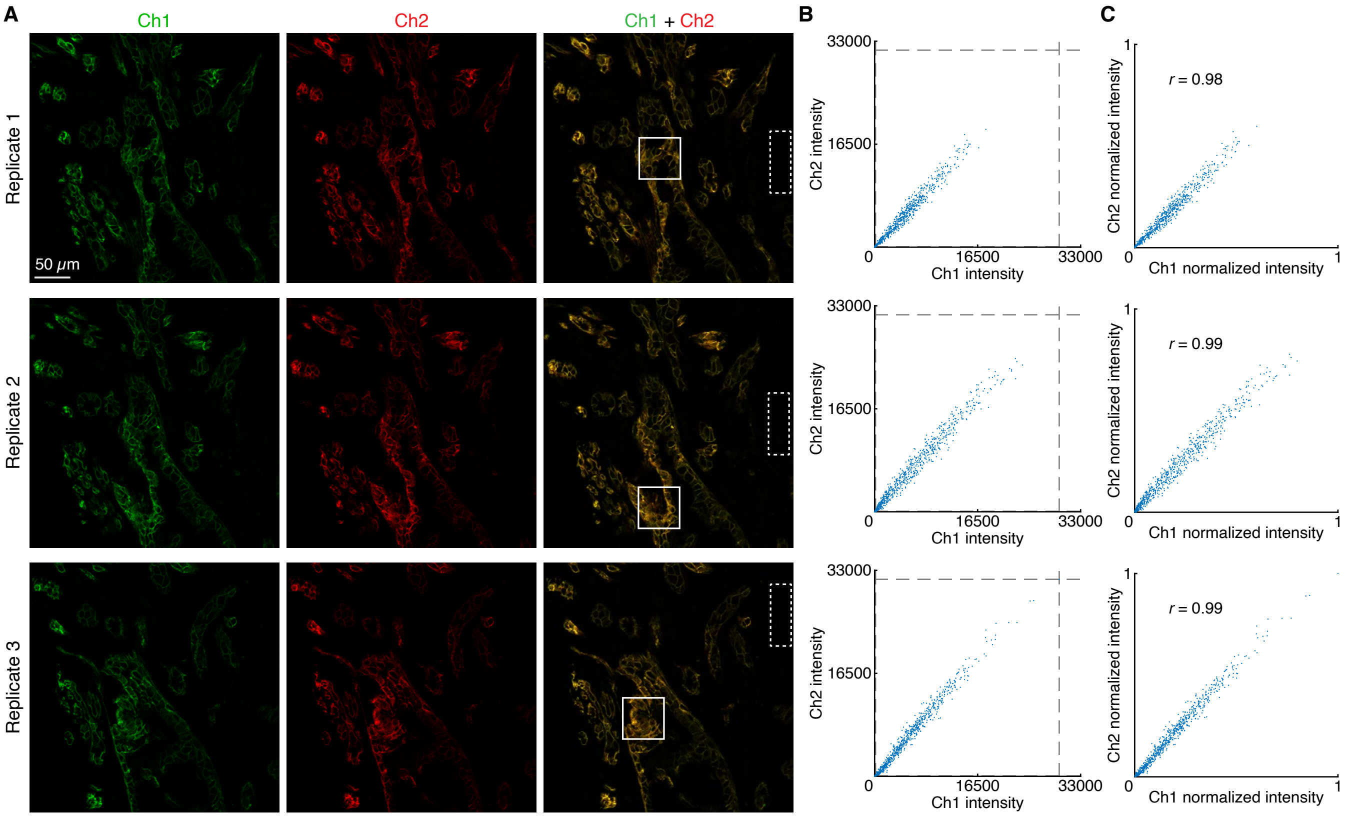

**Figure S17. Redundant 2-channel detection of $\beta$-catenin:E-cadherin target complex in FFPE human breast tissue sections (cf. Figure 5).** (A) Confocal images: individual channels and merge; single optical section. Ch1: $\beta$-catenin:E-cadherin (Alexa546). Ch2: $\beta$-catenin:E-cadherin (Alexa647). Solid boundaries denote regions of variable expression. Dashed boundaries denote regions of no/low expression. Pixel size: $0.568 \times 0.568 \times 3.3$ $\mu$m. Sample: FFPE normal human breast tissue sections. (B) Raw voxel intensity scatter plots representing signal plus background for voxels within solid boundaries of panel A. Voxel size: $2.0 \times 2.0 \times 3.3$ $\mu$m. Dashed lines represent BOT and TOP values (Table S16) used to normalize data for panel C using methods of Section S1.7.3. (C) Normalized voxel intensity scatter plots representing normalized signal (Pearson correlation coefficient, $r$).

| Channel | Protein:protein complex | Fluorophore | BOT | TOP |
|---|---|---|---|---|
| Ch1 | $\beta$-catenin:E-cadherin | Alexa546 | 143 | 29571 |
| Ch2 | $\beta$-catenin:E-cadherin | Alexa647 | 75 | 31564 |

**Table S16. BOT and TOP values used to calculate normalized voxel intensities for scatter plots of Figures 5C (bottom panel) and S17C using methods of Section S1.7.3.** Analysis based on rectangular regions depicted in Figure S17A.

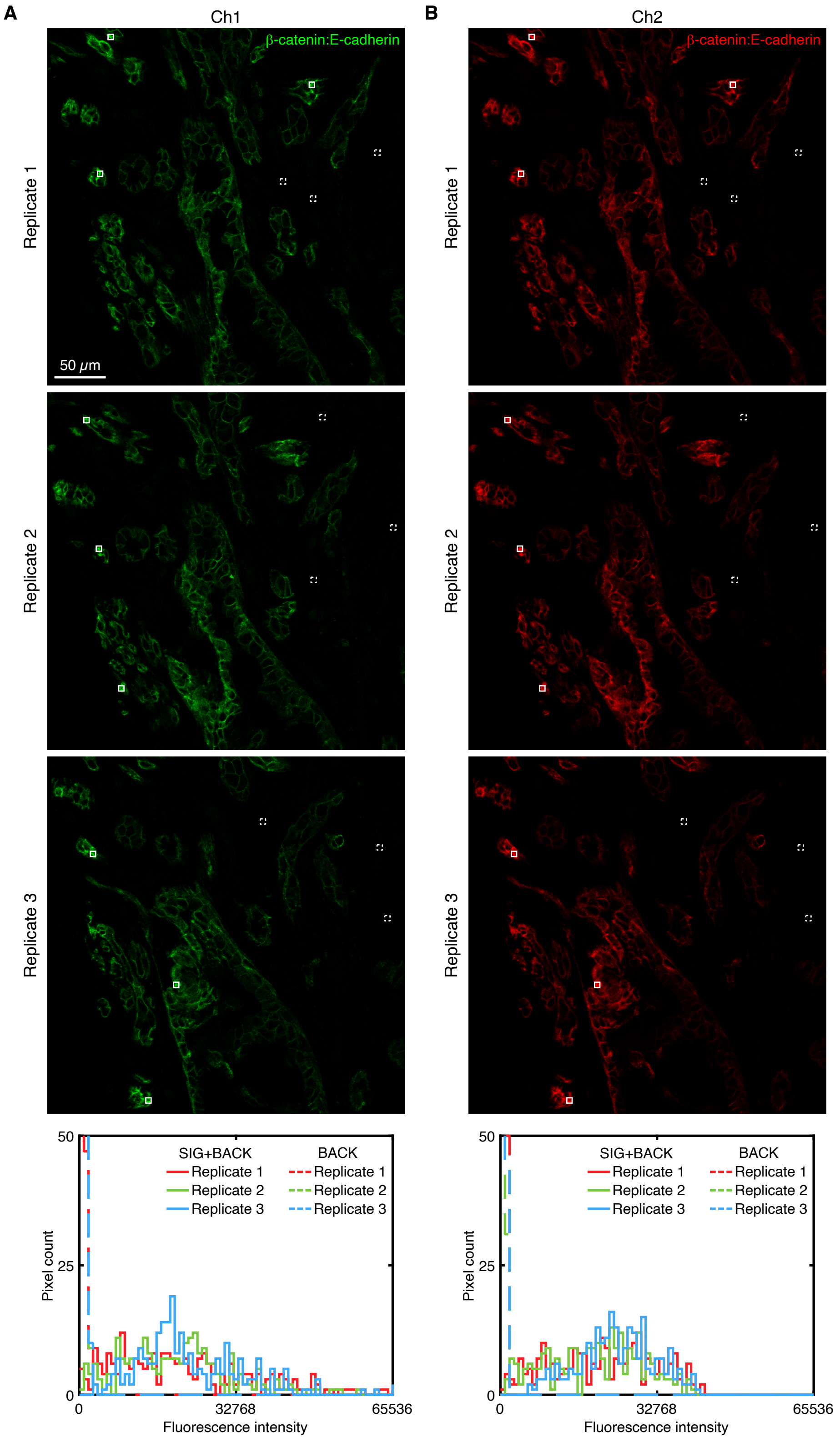

**Figure S18. Measurement of signal and background for redundant 2-channel detection of $\beta$-catenin:E-cadherin target complex in FFPE human breast tissue sections (cf. Figure 5).** (A) Ch1 (Alexa546) of redundant 2-channel imaging of $\beta$-catenin:E-cadherin target complex. (B) Ch2 (Alexa647) of redundant 2-channel imaging of protein:protein complex $\beta$-catenin:E-cadherin. (A,B) Experiment of Type 3 in Table S6. Top: confocal images; single optical section. Solid boundaries denote representative regions of high expression; dashed boundaries denote representative regions of no/low expression. Bottom: pixel intensity histograms for the depicted representative regions. Sample: FFPE normal human breast tissue sections; thickness: 5 $\mu$m.

| Channel | Protein:protein complex | Fluorophore | BACK | SIG+BACK | SIG | SIG/BACK |
|---|---|---|---|---|---|---|
| Ch1 | $\beta$-catenin:E-cadherin | Alexa546 | $410 \pm 50$ | $21\,300 \pm 800$ | $20\,900 \pm 800$ | $52 \pm 6$ |
| Ch2 | $\beta$-catenin:E-cadherin | Alexa647 | $230 \pm 20$ | $21\,000 \pm 1000$ | $21\,000 \pm 1000$ | $91 \pm 9$ |

**Table S17. Estimated signal-to-background for redundant 2-channel detection of $\beta$-catenin:E-cadherin target complex in FFPE human breast tissue sections (cf. Figure 5).** Mean $\pm$ estimated standard error of the mean via uncertainty propagation for $N = 3$ replicate FFPE human breast tissue sections. Analysis based on rectangular regions depicted in Figure S18 using methods of Section S1.7.2.

#### S2.7 Replicates, signal, and background for simultaneous multiplex imaging of protein, protein:protein complex, and RNA targets in human cells (cf. Figure 6)

For simultaneous multiplex imaging of protein, protein:protein, and RNA targets in human cells, the 4 channels are (1 target protein + 1 protein:protein complex + 1 target RNA + DAPI):

- **Ch1:** Target protein HSP60, amplifier B5-Alexa488.
- **Ch2:** Protein:protein complex $\alpha$-tubulin:$\beta$-tubulin, amplifier B1-Alexa546.
- **Ch3:** Target RNA *U6*, amplifier B3-Alexa647.
- **Ch4:** DAPI.

Additional studies are presented as follows:

- Figure S19 displays images for $N = 3$ replicate wells on a multi-well slide (cf. Figure 6).
- Figures S20–S22 display representative regions of individual channels used for measurement of signal and background for each target.
- Table S18 displays estimated values for signal, background, and signal-to-background for each channel.

**Protocol:** Section S1.8.
**Sample:** HeLa cells.
**Reagents:** Table S1.

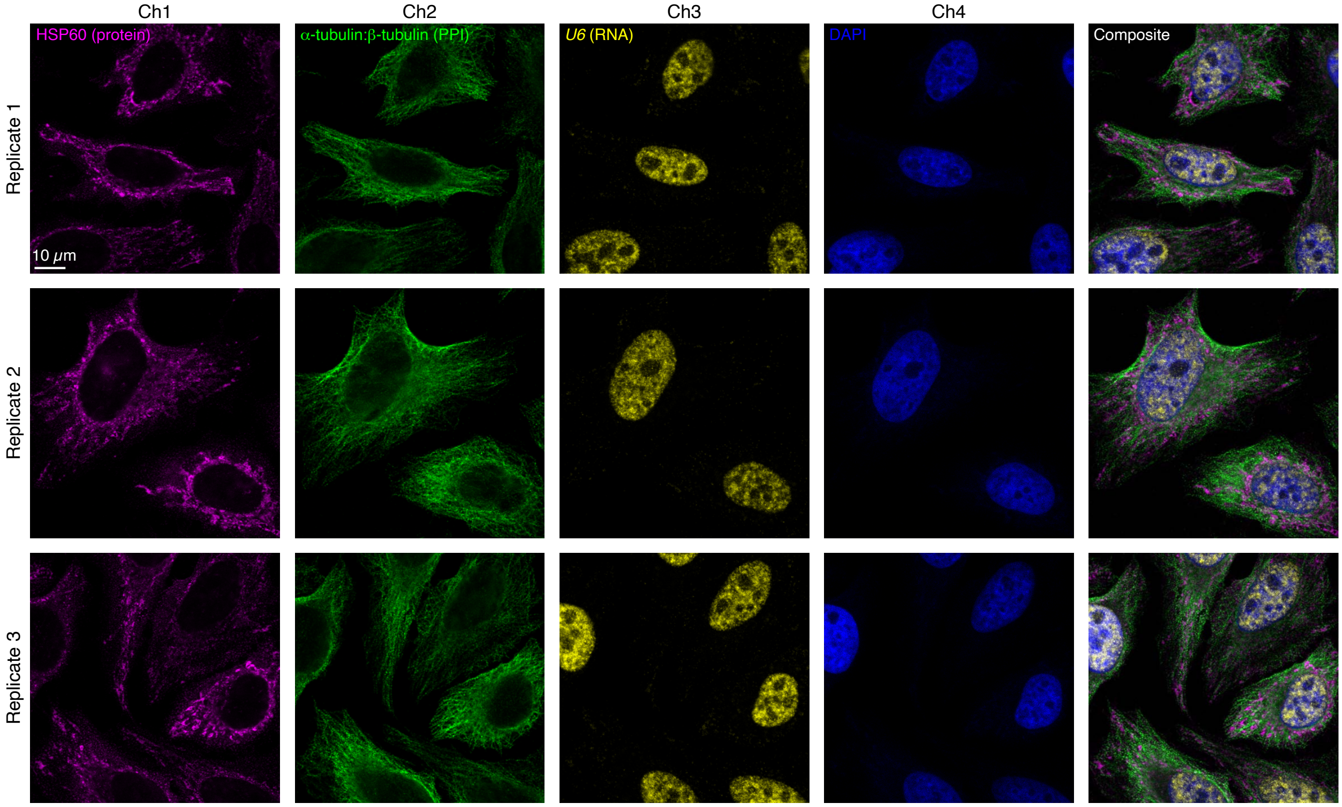

**Figure S19. Replicates for simultaneous multiplex imaging of protein, protein:protein, RNA targets in human cells (cf. Figure 6).** 4-channel images for 3 replicate wells on a multi-well slide; single optical section. Ch1: target protein HSP60 (Alexa488). Ch2: protein:protein complex $\alpha$-tubulin:$\beta$-tubulin (Alexa546). Ch3: target RNA *U6* (Alexa647). Ch4: DAPI. Sample: HeLa cells.

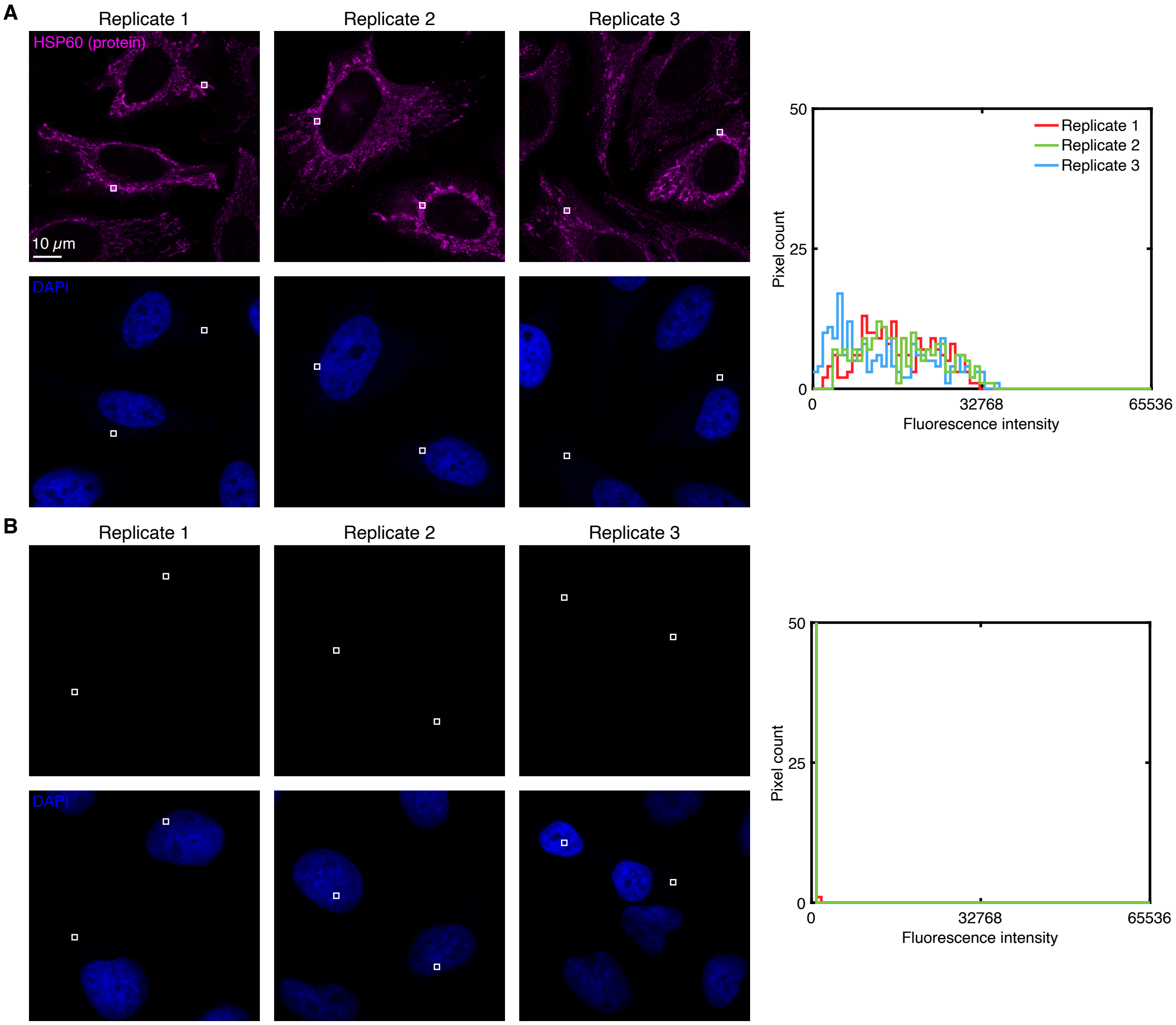

**Figure S20. Measurement of signal and background for target protein HSP60 using HCR IHC in human cells (cf. Figure 6).** (A) Use experiment of Type 1 in Table S7 to measure SIG+BACK in a region of high target protein expression (pixels within rectangles) in 3 replicate wells. (B) Use experiment of Type 2 in Table S7 to measure BACK $\approx$ $NSD_{2^\circ}$+NSA+AF in a region of maximum background (pixels within rectangles) in 3 replicate wells. (A,B) Left: individual channels of confocal images collected with the microscope settings optimized to avoid saturating SIG+BACK pixels; DAPI channel contextualizes placement of the rectangles; single optical section. Ch1: target protein HSP60 (Alexa488). Ch4: DAPI. Right: pixel intensity histograms for rectangular regions of Ch1 (one rectangle in each of 2 individual cells in each of 3 replicate wells on a multi-well slide).

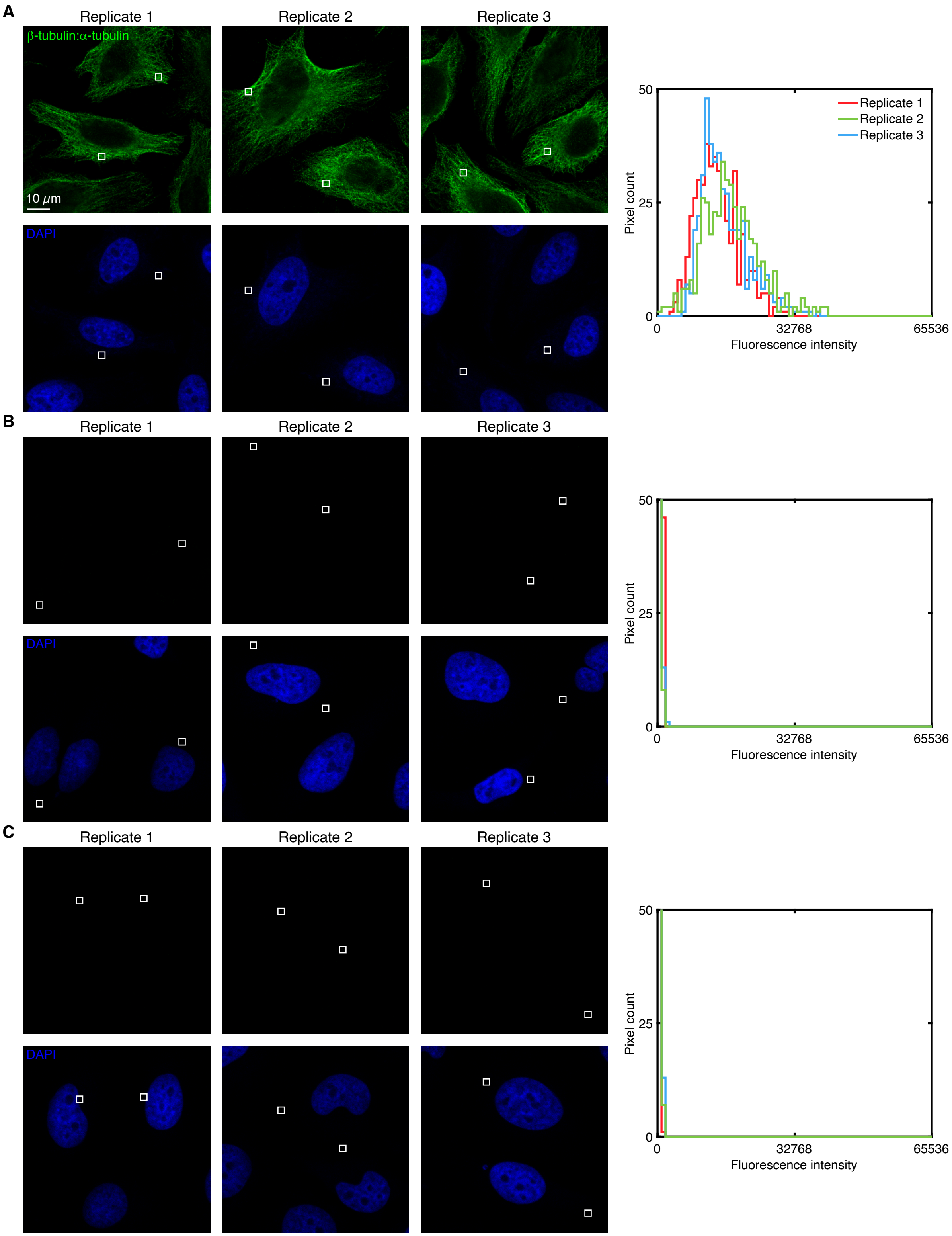

**Figure S21. Measurement of signal and background for protein:protein complex $\beta$-tubulin:$\alpha$-tubulin in human cells (cf. Figure 6).** (A) Use experiment of Type 1a in Table S6 to measure SIG+BACK in a region of high protein:protein complex expression (pixels within rectangles) in 3 replicate wells. (B) Use experiment of Type 2a in Table S6 to measure $NSD_{P1}$+NSA+AF in a region of maximum background (pixels within rectangles) in 3 replicate wells. (C) Use experiment of Type 2b in Table S6 to measure $NSD_{P2}$+NSA+AF in a region of maximum background (pixels within rectangles) in 3 replicates. (B,C) Background estimated as BACK $\approx$ $NSD_{P1}$+NSA+AF + $NSD_{P2}$+NSA+AF. (A,B,C) Left: individual channels of confocal images collected with the microscope settings optimized to avoid saturating SIG+BACK pixels; DAPI channel contextualizes placement of the rectangles; single optical section. Ch2: protein:protein complex $\alpha$-tubulin:$\beta$-tubulin (Alexa546). Ch4: DAPI. Right: pixel intensity histograms for rectangular regions of Ch2 (one rectangle in each of 2 individual cells in each of 3 replicate wells on a multi-well slide).

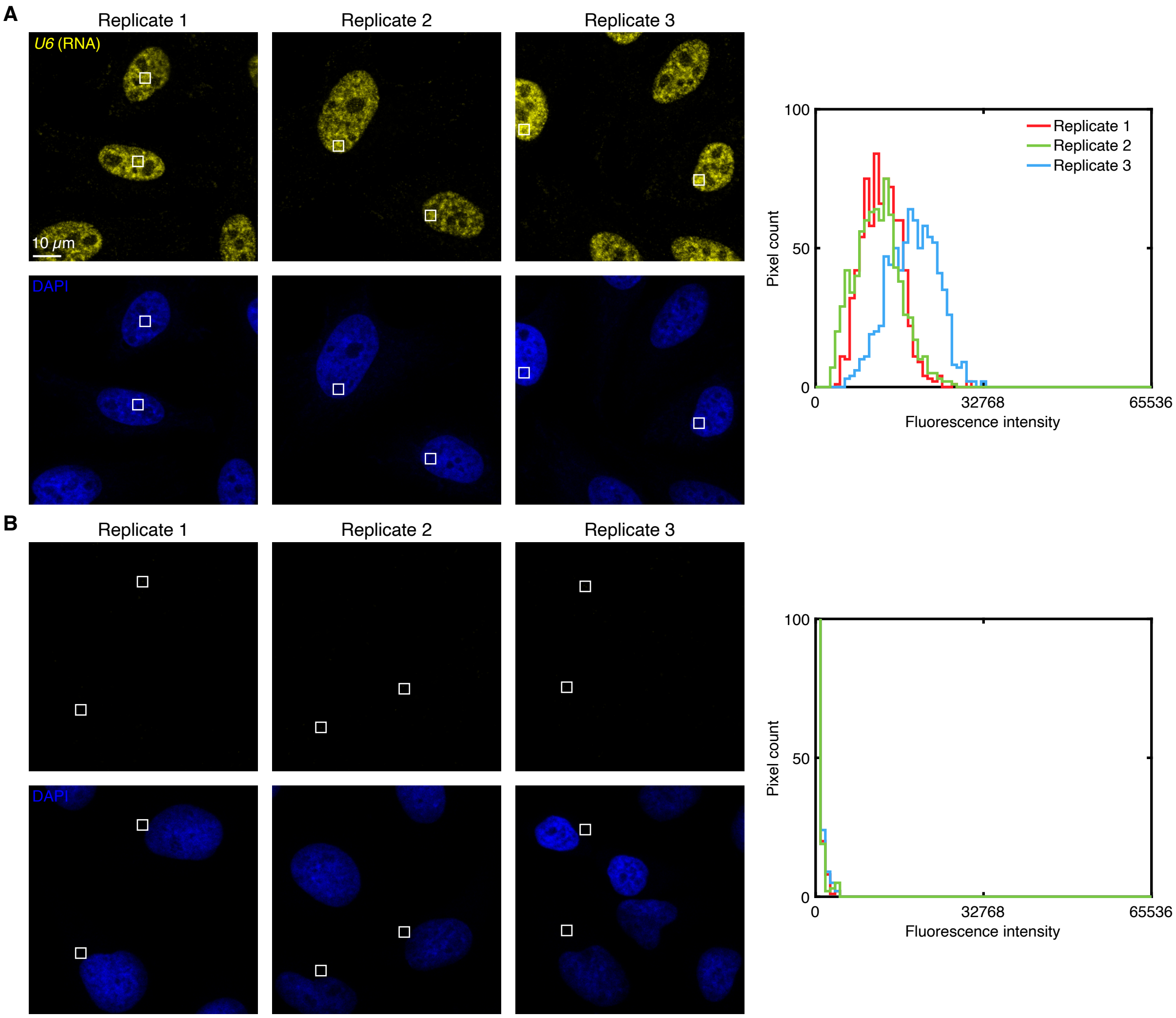

**Figure S22. Measurement of signal and background for target RNA *U6* in human cells (cf. Figure 6).** (A) Use experiment of Type 1 in Table S8 to measure SIG+BACK in a region of high target RNA expression (pixels within rectangles) in 3 replicate wells. (B) Use experiment of Type 2 in Table S8 to measure BACK≈ NSA+AF in a region of maximum background (pixels within rectangles) in 3 replicate wells. (A,B) Left: individual channels of confocal images collected with the microscope settings optimized to avoid saturating SIG+BACK pixels; DAPI channel contextualizes placement of the rectangles; single optical section. Ch3: target RNA *U6* (Alexa647). Ch4: DAPI. Right: pixel intensity histograms for rectangular regions of Ch3 (one rectangle in each of 2 individual cells in each of 3 replicate wells on a multi-well slide).

| Target or complex | SIG+BACK | $NSD_{P1}$+NSA+AF | $NSD_{P2}$+NSA+AF | BACK | SIG | SIG/BACK |
|---|---|---|---|---|---|---|
| HSP60 | $15\,000 \pm 1000$ | — | — | $21 \pm 3$ | $15\,000 \pm 1000$ | $800 \pm 100$ |
| $\alpha$-tubulin:$\beta$-tubulin | $15\,500 \pm 900$ | $90 \pm 40$ | $54 \pm 3$ | $140 \pm 40$ | $15\,400 \pm 900$ | $110 \pm 30$ |
| *U6* | $14\,000 \pm 2000$ | — | — | $90 \pm 6$ | $14\,000 \pm 2000$ | $160 \pm 30$ |

**Table S18. Estimated signal-to-background for simultaneous multiplex imaging of protein, protein:protein, and RNA targets in human cells (cf. Figure 6).** Mean $\pm$ estimated standard error of the mean via uncertainty propagation for representative regions of $N = 3$ replicate wells on a multi-well slide. Analysis based on rectangular regions depicted in Figures S20–S22 using methods of Section S1.7.2.
